# Spatiotemporal DNA Methylome Dynamics of the Developing Mammalian Fetus

**DOI:** 10.1101/166744

**Authors:** Yupeng He, Manoj Hariharan, David U. Gorkin, Diane E. Dickel, Chongyuan Luo, Rosa G. Castanon, Joseph R. Nery, Ah Young Lee, Brian A. Williams, Diane Trout, Henry Amrhein, Rongxin Fang, Huaming Chen, Bin Li, Axel Visel, Len A. Pennacchio, Bing Ren, Joseph R. Ecker

## Abstract

Genetic studies have revealed an essential role for cytosine DNA methylation in mammalian development. However, its spatiotemporal distribution in the developing embryo remains obscure. Here, we profiled the methylome landscapes of 12 mouse tissues/organs at 8 developmental stages spanning from early embryogenesis to birth. Indepth analysis of these spatiotemporal epigenome maps systematically delineated ~2 million methylation variant regions and uncovered widespread methylation dynamics at nearly one-half million tissue-specific enhancers, whose human counterparts were enriched for variants involved in genetic diseases. Strikingly, these predicted regulatory elements predominantly lose CG methylation during fetal development, whereas the trend is reversed after birth. Accumulation of non-CG methylation within gene bodies of key developmental transcription factors coincided with their transcriptional repression during later stages of fetal development. These spatiotemporal epigenomic maps provide a valuable resource for studying gene regulation during mammalian tissue/organ progression and for pinpointing regulatory elements involved in human developmental diseases.

## Introduction

Mammalian embryonic development involves spatiotemporal transcriptional regulation, which is mediated by sophisticated orchestration of epigenetic modifications and transcription factor (TF) binding of regulatory DNA elements, primarily enhancers. The accessibility of TFs to regulatory DNA is closely related to the covalent modification of histones and DNA^1–6^.

Cytosine DNA methylation (mC) is an epigenetic modification that plays a critical role in gene regulation^7^. In the mammalian genome, mC occurs predominantly on cytosine followed by guanine (mCG), which is dynamic at regulatory elements in different tissues and cell types^8–12^. In fact, mCG is able to directly affect the DNA binding affinity of a variety of transcription factors^1,2,6,13–15^ and targeted removal/addition of mCG in promoters is concomitant with increases/decreases in gene transcription^16^. Non-CG methylation (mCH; H = A, C or T) is also present at an appreciable level in embryonic stem cells, oocytes as well as brain, heart, skeletal muscle and a variety of adult tissues^8–10,17–20^. Although its precise function(s) are unknown, mCH abundance directly affects DNA binding of MeCP2, the methyl-binding protein responsible for Rett Syndrome^20–23^.

mC is actively regulated during mammalian development^24,25^. However, in contrast to pre-implantation embryogenesis^25–27^, data are lacking for the later stages of fetal development, during which anatomical features of the major organ systems are more evident and human birth defects are manifested^28^. To fill this knowledge gap, we used the experimentally tractable mouse embryo as a model system to profile DNA methylation variation. Deep whole-genome bisulfite sequencing^8^ was performed to comprehensively profile cytosine DNA methylation in 12 tissue types (in replicate) for 8 development stages starting from embryonic day 10.5 (E10.5) to birth (postnatal day 0, P0; Fig. 1A). The temporal mouse fetal tissue/organ epigenomes (Supplemental Table 1) described here should inform our understanding of regulatory events occurring during normal human fetal progression as well as developmental disorders. These comprehensive datasets are publically accessible at https://www.encodeproject.org/search/?searchTerm=ecker&type=Experiment&award.rfa=ENCODE3 and http://neomorph.salk.edu/ENCODE_mouse_fetal_development/.

**Figure 1.**
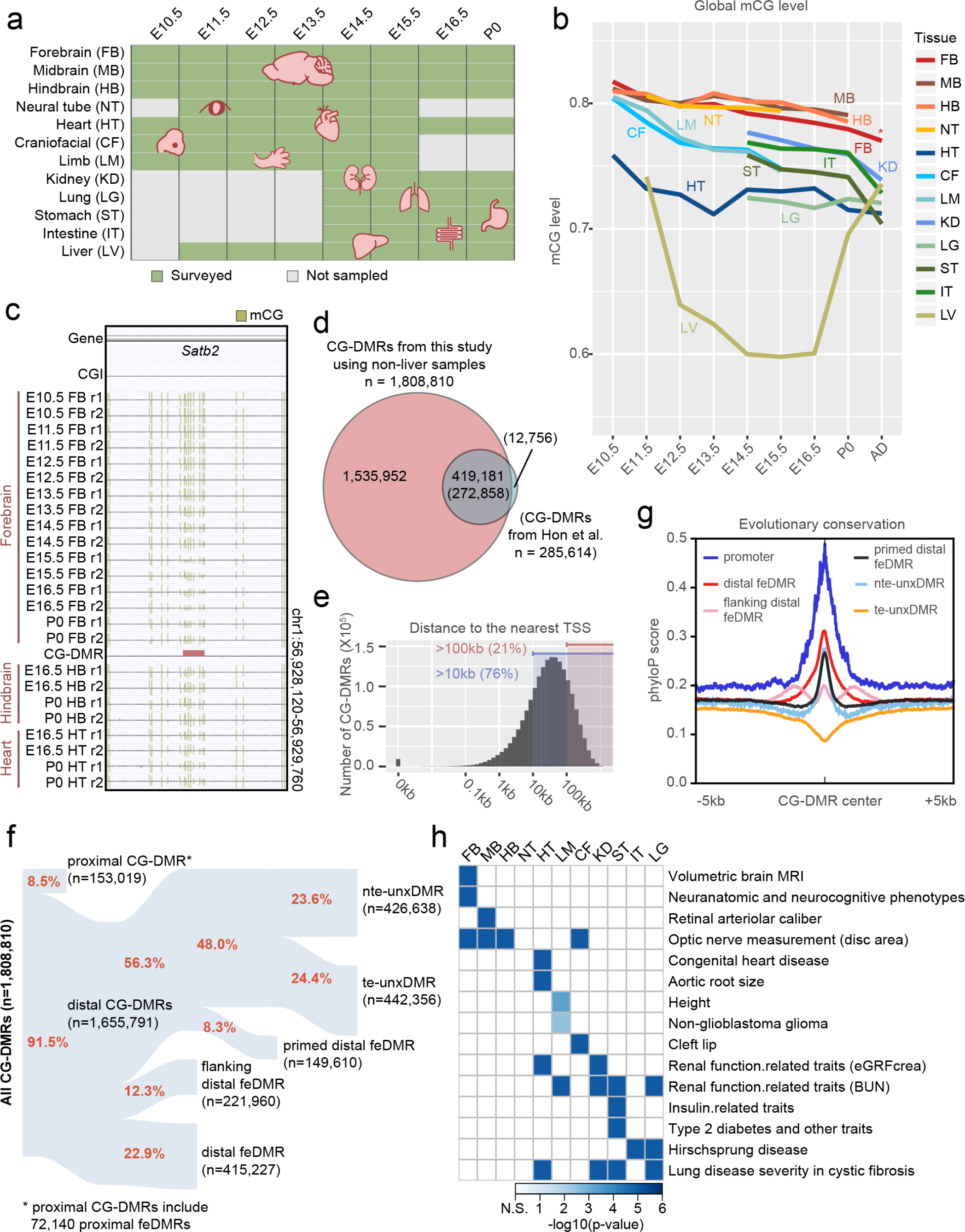
Annotation of methylation variable regulatory elements in developing mouse tissues. **a**, Tissue samples (green) profiled in this study. Grey cells mark the tissues and stages that were not sampled because either the tissue is not yet formed or it was not possible to obtain enough material for the experiment, or the tissue-type was too heterogeneous to obtain informative data. **b**, Genome-wide CG methylation (mCG) levels of each tissue across their developmental trajectories. The data point of adult forebrain (postnatal 6 week frontal cortex) is from Lister et al^10^. **c**, Example of a CG differentially methylated region (CG-DMR) in the body of the *Satb2* gene. Top two tracks show the gene annotation and the locations of CG islands (CGIs), which are followed by mCG tracks and a CG-DMR track. Gold ticks represent methylated CG sites and their heights indicate the mCG level, ranging from 0 to 1. Ticks on the forward DNA strand are projected upward and ticks on the reverse DNA strand are projected downward. **d**, Fetal CG-DMRs identified in this study encompass the majority of the adults CG-DMRs from a previous study of adult tissues^6^. The numbers related to fetal CG-DMR in this study are shown without parenthesis, whereas the numbers in parenthesis are related to adult tissue CG-DMRs. **e**, Distance of CG-DMRs to the nearest transcription start sites (TSSs). **f**, Categorization of CG-DMRs. “proximal CG-DMRs” are CG-DMRs that are overlapped with promoters, CG islands (CGIs) or CGI shores. The remaining CG-DMRs are defined as “distal CG-DMRs”. fetal enhancer-linked CG-DMRs (feDMRs) are those predicted to show enhancer activity using REPTILE algorithm^27^, which contain 415,227 distal feDMRs and 72,140 proximal feDMRs. CG-DMRs within 1kb to distal feDMRs are “flanking distal CG-DMRs”. Of the remaining distal CG-DMRs, we defined “primed distal feDMRs” as those showing strong hypomethylation patterns (mCG difference >= 0.3) and also primed enhancer-like chromatin signatures. The remaining CG-DMRs are unexplained distal CG-DMRs” (unxDMRs), whose functions are unknown. unxDMRs are further stratified based on their overlap with transposons: transposal element overlapping unxDMRs (“te-unxDMRs”) and transposal element non-overlapping unxDMRs (“nte-unxDMRs”). The number of CG-DMRs assigned to each group is shown in the parentheses. See Methods for details. **g**, Conservation (phyloP) score of promoters and different categories of distal CG-DMRs. **h**, Tissue-specific feDMRs are enriched in GWAS SNPs associated with tissue/organ specific functions and tissue-related disease states.

## Results

### CG methylation dynamics

To assess the mC landscape in the developing mouse embryo, 144 high-quality tissue methylomes were produced that cover most of the major organ systems and tissue types derived from the three primordial germ layers. To obtain an overview of the spatiotemporal profile of fetal tissue/organ methylome reconfiguration, we first calculated the global mCG level at each developmental stage (Fig. 1a-b; Methods). The genomes of all fetal tissues were heavily CG methylated, ranging from 70%-82% with the notable exception of liver, which ranged from 60%-74% of CGs and showed large domains with partial methylation. Interestingly, such large partially methylated domains (PMDs), a genomic feature previously observed in only human cultured cell lines^8,29^, pancreas^9^, placenta^30^ and cancer samples^31,32^ were found exclusively in mouse fetal liver (Extended Data Fig. 1). PMD formation and dissolution precisely coincided with fetal liver hematopoiesis (see Supplemental Note). Finally, tissues primarily derived from ectoderm (forebrain, midbrain, hindbrain and neural tube) showed overall higher mCG levels than other tissue types (heart, craniofacial, limb, kidney, lung, stomach, intestine and liver).

Despite similar global mCG levels in fetal tissues, we identified 1,808,810 CG differentially methylated regions (CG-DMRs), which are on average 339bp long and cover 22.5% (614 Mb) of the mouse genome ((Fig. 1c; Methods). Comprehensive fetal tissue CG-DMR annotation captured ~96% (n=272,858) of all previously reported adult mouse tissue CG-DMRs^12^, while finding over 1.5 million new regions (Fig. 1d). Surprisingly, only a minority (8.5% or 153,019) of CG-DMRs overlapped with promoters, CpG islands (CGIs) or CGI shores (Fig. 1e-f; Extended Data Fig. 2a-c; Methods). The vast majority of CG-DMRs (~92.5% or 1,655,791) were distally located and showed a high degree of evolutionary conservation, suggesting that they are functional (Fig. 1f-g).

### Annotating methylation variable regions

To further classify fetal CG-DMRs, we delineated those genomic regions likely associated with enhancer activity using the REPTILE^33^ algorithm which allows enhancer prediction by integration of the fetal tissue mCG data and histone modification chromatin immunoprecipitation sequencing (ChIP-seq) data (Gorkin et al companion paper, this issue; Supplemental Table 2). Considering all fetal tissues, except liver, we identified 487,367 CG-DMRs as fetal enhancer-linked CG-DMRs or feDMRs, 85% (415,227) of which are distal to known promoters (Fig. 1f; Methods). feDMRs are evolutionarily conserved and show enhancer-like chromatin signatures including the depletion of mCG and H3K27me3, and the enrichment of H3K4me1 and H3K27ac^8,34–36^ (Fig. 1g; Extended Data Fig. 2d). Many (106,016) of these enhancers were not previously reported in adult mouse tissues^37^ (Extended Data Fig. 3a). More than 60% of the feDMRs-overlapping VISTA enhancer elements^38^ showed *in vivo* enhancer activity in E11.5 mouse embryo, and this percentage increased for higher scoring feDMRs (Extended Data Fig. 3b). Moreover, tissue-specific feDMRs are enriched for TF binding motifs related to specific tissue function(s) and are near genes in specific tissue-related pathways (Extended Data Fig. 3c; Supplemental Table 3). We found that the human orthologous regions of feDMRs significantly overlapped with the disease/trait-associated single nucleotide polymorphisms identified from genome-wide association studies and showed tissue-specificity (Fig. 1h; Supplemental Table 4; Methods).

We also identified 221,960 CG-DMRs that flank (within 1kb) feDMRs but which are not themselves predicted as enhancers (Fig. 1g). These flanking distal feDMRs (fd-feDMRs) are much less conserved than feDMRs although their mCG level is moderately correlated with nearby feDMRs (mean Pearson correlation coefficient = 0.41; see Methods), suggesting that a fraction of fd-feDMRs may be the by-product of demethylation of the adjacent feDMRs. Alternatively, fd-feDMRs may be bound by pioneer TF(s) which allow opening of chromatin of adjacent feDMRs^39,40^. This notion is supported by the enrichment of binding motifs for several known pioneer TFs^39^ such as FOXA2, GATA3 and PBX1 at fd-DMRs (Supplemental Table 5). Interestingly, binding motifs of the insulator protein CTCF and several transcriptional repressors (e.g. CUX1^41^) are also enriched at fd-feDMRs, indicating a third possibility that fd-feDMRs correspond to insulator and silencer elements (Supplemental Table 5).

Besides the above CG-DMR classes, another type of distal CG-DMR (n = 149,610) are the primed distal feDMRs (pd-feDMRs); these display strong CG hypomethylation in at least one tissue sample and are linked to primed fetal enhancers^42^ (mCG difference ≥ 0.3; Extended Data Fig. 4a; Methods). In the tissues where they are hypomethylated, pd-feDMRs showed chromatin signatures resembling primed enhancers^42^ which are enriched for H3K4me1 while lacking H3K27ac and H3K27me3 (Extended Data Fig. 4b). Like feDMRs, pd-feDMRs are also evolutionary conserved (Fig. 1g). Consistent with their putative role as enhancers, and similar to feDMRs, they share significant TF-binding motif signature enrichment in 9 out of the 12 tissue types (Extended Data Fig. 4c; Supplemental Table 6).

The remaining unclassified distal CG-DMRs (868,994) show more subtle CG hypomethylation patterns, suggesting that they likely derive from a small fraction of cells within these complex tissues/organs (Methods). Since a functional role cannot yet be assigned, we named this group “unexplained CG-DMRs” (unxDMR). The genomic locations of unxDMRs significantly overlapped with transposable elements (TEs; 58.6%, p-value < 0.001; Methods). Inspired by this observation, we divided unxDMRs into two subgroups: unxDMRs that overlapped with TE (te-unxDMRs) and ones not overlapping (nte-unxDMRs) (Fig. 1F; Extended Data Fig. 4D and E). te-unxDMRs were less evolutionarily conserved compared to flanking regions but transposons may be a source of novel regulatory elements^43^. Different from te-unxDMRs, genomic sequences underling nte-unxDMRs are as conserved as feDMRs, implying that they may be functional. Indeed, 10%-45% of the nte-unxDMRs showed open chromatin in purified neurons^44^ and/or a variety of mouse cell lines and tissues^45^ (p-value < 1e-3; permutation test; Methods). Therefore, nte-unxDMRs are likely regulatory elements active only in rare cell types and their weak hypomethylation profiles are due to the tissue heterogeneity, highlighting the future necessity of cell type-specific^46^ or single-cell epigenomic studies^47^.

### Distinct pre- and postnatal mCG dynamics

The dominant methylation pattern that emerged during fetal progression was a continuous loss-of-mCG at tissue-specific CG-DMRs, which strongly overlap with predicted enhancers (Fig. 2a; Extended Data Fig. 5a-b; Methods). In striking contrast, the gain-of-mCG methylation at CG-DMRs mainly occurred after birth (Fig. 2a; Extended Data Fig. 5a). To quantify these changes for each stage interval, we counted loss-of-mCG (mCG decreasing by at least 0.1 in one CG-DMR) and gain-of-mCG events (mCG increasing by at least 0.1 in one CG-DMR) (Fig. 2b; Methods). During the period from E10.5 to P0, 77% to 95% of the mCG changes involved loss-of-mCG, depending on the tissue examined. More than 70% of the loss-of-mCG events occurred between E10.5 to E13.5 in all tissues except in heart (44%; Fig. 2a; Extended Data Fig. 5c). The mCG level of 44-84% tissue-specific CG-DMRs dropped to below 0.5 at E14.5, compared to only 16-31% at E10.5. Since allele-specific methylation is relatively rare^9^, the observed methylation dynamics suggest that after E14.5, most of the tissue-specific CG-DMRs are unmethylated in more than half of the cells in a tissue.

**Figure 2.**
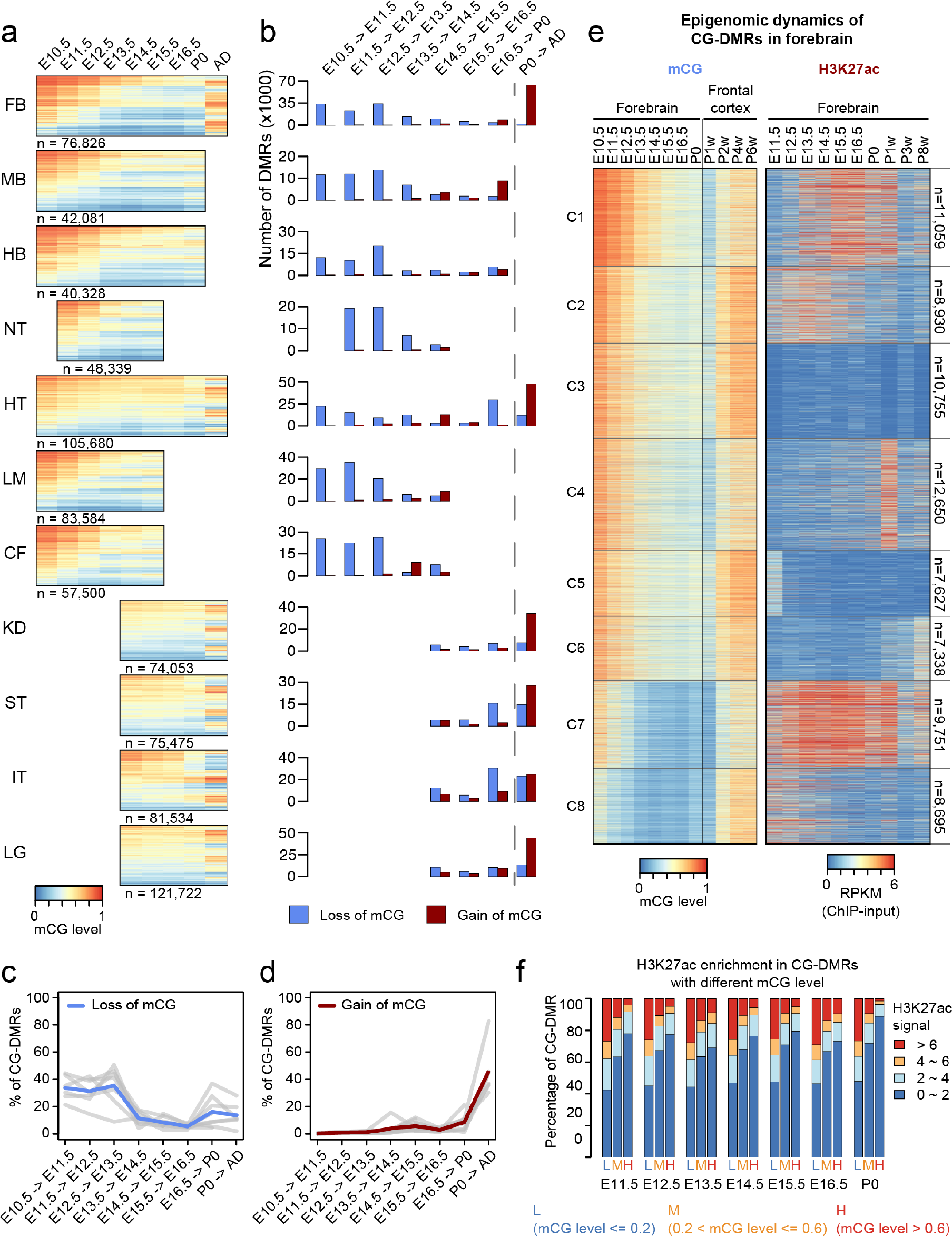
Tissue-specific CG-DMRs undergo continuous demethylation during embryogenesis and remethylation after birth. **a**, CG methylation (mCG) level of tissue-specific CG-DMRs. The number under each heatmap indicates the number of tissue-specific CG-DMRs. mCG data from adult (AD) forebrain was approximated using data from postnatal 6 week frontal cortex from Lister et al^10^. **b**, The numbers of loss-of-mCG events (blue) and gain-of-mCG events (red) in tissue-specific CG-DMRs for each fetal stage interval. We defined one loss-of-mCG (gain-of-mCG) event as the decrease (increase) of at least 0.1 in mCG level of one CG-DMR for one fetal stage interval. **c-d**, Fraction of tissue-specific CG-DMRs that undergo lost-of-mCG (blue) and gain-of-mCG (red) during development. The blue (loss-of-mCG) or red (gain-of-mCG) line shows the aggregated values over all non-liver tissues, whereas grey lines show the data for each tissue type. **e**, mCG and H3K27ac dynamics of forebrain-specific CG-DMRs. Frontal cortex methylomes from postnatal 1, 2, 4, 6 weeks (P1w to P6w) were compared with data from adult forebrain. Forebrain-specific CG-DMRs were clustered into 8 groups (see Methods for details). **f**, Relationship between mCG level and enrichment of H3K27ac in tissue-specific CG-DMRs. For each tissue type, tissue-specific CG-DMRs were first grouped into three categories (L: low; M: median; H: high) based on their mCG level. Then, the fraction of tissue-specific CG-DMRs from each category that showed different levels of H3K27ac enrichment was quantified. This panel shows the results of all non-liver tissues.

Compared to the loss of mCG, the vast majority (57%-86%) of gain-of-mCG events happened after birth (Extended Data Fig. 5d). As a result, 27%-56% of the tissue-specific CG-DMRs become highly methylated (mCG level > 0.6) in adult tissues (at least 4 weeks old), while the number is 0.3%-15% at birth (P0), reflecting the silencing of fetal regulatory elements (Extended Data Fig. 6a). The absence of gain-of-mCG events during fetal development may be a result of limited *de novo* CG methylation and/or excessive demethylation activity.

The observed mCG dynamics cannot be explained by the lack of expression of cytosines methytransferases *Dnmt1* and *Dnmt3a* nor the co-factor *Uhrf1*^48^, all of which are highly expressed between E10.5 and E13.5 when major the loss-of-mCG events occur (Extended Data Fig. 6b). Furthermore, we also did not find increased expression of *Tet* methylcytosine dioxygenases, involved methylation removal^49^, during the same period. Interestingly, *Tet3* expression is lower in heart, coinciding with its less dynamic mCG during embryogenesis. Absence of gain-of-methylation events until the postnatal period may involve translational and/or posttranslational regulation of these enzymes or possibly other unknown control mechanisms. Importantly, WGBS does not distinguish between 5-methylcytosine and 5-hydroxymethylcytosine^50^. Based on earlier studies^10,51^, the contribution of 5-hydroxymethylcytosine to the total of cytosine modifications can be relatively minor. However, future studies that directly measure the full complement of cytosine modifications (5-hydroxymethylcytosine, 5-formylcytosine and 5-carboxylcytosine) are needed to understand their dynamics during fetal tissue development.

### Dynamic mCG and chromatin states

To further pinpoint the timing of CG-DMR remethylation and its relationship with enhancer activity, we included methylation data from adult frontal cortex^10^ as well as H3K27ac ChIP-seq data from adult forebrain^52^ (Supplemental Table 2). Using these additional datasets, we further refined the timing of remethylation in the frontal cortex. We clustered forebrain-specific CG-DMRs into 8 groups based on the mCG and H3K27ac dynamics across both fetal and adult stages (Fig. 2e; Extended Data Fig. 6c; Methods). Between the 1^st^ and 2^nd^ postnatal weeks, methylation of forebrain-specific CG-DMRs increased dramatically and was even further methylated during tissue maturation (Extended Data Fig. 6d).

We then asked how the mCG dynamics at tissue-specific CG-DMRs were associated with their enhancer activity (approximated by H3K27ac abundance) during development. Interestingly, although the temporal depletion of mCG was not necessarily related to high H3K27ac enrichment (e.g. cluster 3, 5 and 6), a high cytosine methylation state was indicative of a low H3K27ac state (Fig. 2e-f). Only 2-9% of the highly methylated CG-DMRs (mCG level > 0.6) showed high H3K27ac enrichment (>6) and the numbers increase to 25-28% for lowly methylated CG-DMRs (mCG level < 0.2; Fig. 2f). These observations suggest that the decreasing methylation during development may prime dynamic regulation of enhancer activity promoted by increased TF binding and/or altered histone modifications.

### mCG dynamics and gene module expression

We next investigated the association between differential mCG and the transcription of genes in different pathways, using RNA-seq data from the same tissue/organ samples (Wold et al. see companion paper, this issue). By applying an unsupervised, weighted correlation network analysis (WGCNA)^53^ method, we identified 33 co-expressed gene clusters (co-expression modules, CEMs) and calculated “eigengenes” to summarize the expression profile of genes within modules (Fig. 3a-b; Extended Data Fig. 7a-b; Methods). Genes sharing similar expression profiles are more likely to be regulated by a common mechanism and/or involved in the same pathway. For example, CEM12 contains genes that are highly expressed in early developmental stages but are down regulated as tissues mature (Fig. 3c; Extended Data Fig. 7b). Genes in CEM12 are significantly related to cell cycle, matching our knowledge that cells become post-mitotic in mature tissues. Similarly, genes in CEM3 are related to chromatin modification and are lowly expressed in heart, which showed less mCG dynamics relative to other tissues (Fig. 3b-c; Extended Data Fig. 7c). Overall, genes in CEMs are associated with different pathways and/or biological processes (Extended Data Fig. 7d; Supplemental Table 7).

**Figure 3.**
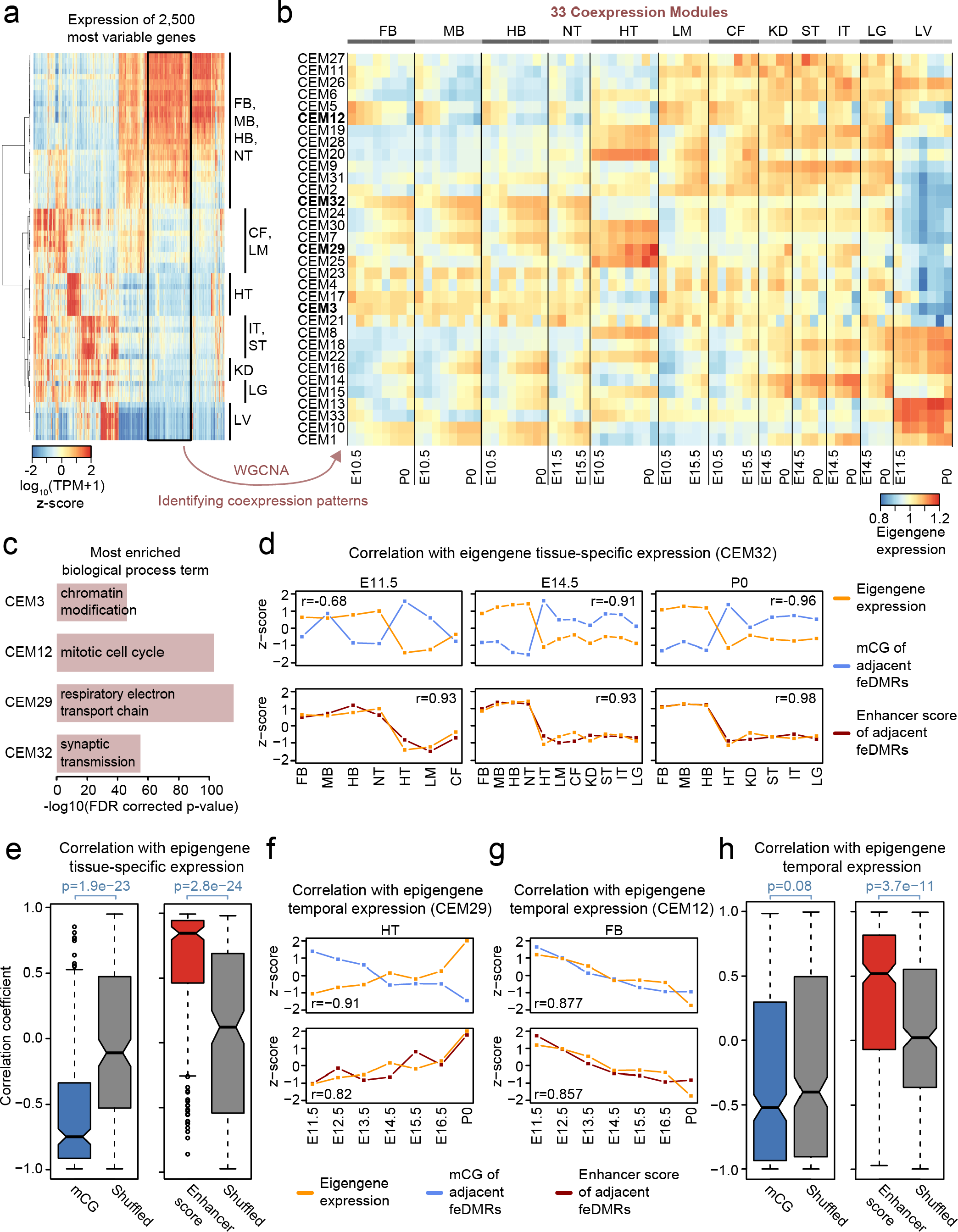
Association between CG methylation (mCG) and the expression patterns of coexpression modules (CEMs). **a**, Expression of the 2,500 most variable genes in all tissue samples. Tissue samples were grouped using hierarchical clustering. Gene expression is measured by log_10_ (TPM+1) (transcripts per million) z-score. Black box highlights a group of co-expressed genes that are highly expressed in neuronal tissues. **b**, 33 CEMs identified in WGCNA and their eigengene expression. The CEMs related to the next panel (c) are bolded. **c**, The most enriched gene ontology (Biological Process) terms of genes in four representative modules. **d**, Correlation of the tissue-specific eigengene expression (orange) for each developmental stage with the average mCG level (blue) and the average enhancer score (red) for feDMRs linked to the genes in the CEM32. The mCG levels (enhancer score) of each feDMR was normalized by dividing the genome-wide average mCG level (enhancer score) and then transformed to z-scores (See Methods for details). Pearson correlation coefficient (r) was calculated. **e**, Pearson correlation coefficients of mCG (blue) or enhancer score (red) of neighboring feDMRs with tissue-specific eigengene expression across all 33 CEMs on all stages. P-values were obtained by testing the median of the Pearson correlation coefficients against the median of the shuffled (grey) using two-tailed Mann-Whitney test. **f-g**, Similar to (d), correlation of temporal eigengene expression for CEM29 (f) and CEM12 (g) with the average mCG level and the average enhancer score of neighboring feDMRs. **h**, Similar to (e), Pearson correlation coefficients of mCG and enhancer score with temporal epigengene expression across all CEMs and all tissue types, excluding liver. See Methods for details.

To understand how mCG profiles and enhancer activity of regulatory elements (feDMRs) are associated with the expression of genes in CEMs, we first linked feDMRs to the neighboring genes. Next, for each CEM, we correlated its eigengene expression with the average mCG and enhancer score (as approximate enhancer activity) of the feDMRs linked to the genes in that CEM (Methods). To tease out the effect of tissue type and development, we calculated the correlations separately for tissue-specific expression and temporal expression (Methods). These analyses revealed that for a given developmental stage methylation of feDMRs was negatively correlated with the tissue-specific eigengene expression, consistent with its known repressive role (Fig. 3d-e). In contrast, the enhancer score of feDMRs was positively correlated with tissue-specific transcription, implying that enhancers likely drive tissue-specific expression (Fig. 3d-e). Such trends hold for the correlations across all modules (Fig. 3e; Methods). Next, for a given tissue type, we calculated the correlation across developmental time. The enhancer score remains positively correlated with the temporal expression, implying that enhancer activity is also the driver of temporal gene expression (Fig. 3f-g). In contrast, because mCG generally decreased at feDMRs over development, it only showed marginally better anti-correlation than that expected by chance alone (Fig. 2a; Fig. 3f-g).

### Large tissue-specific CG-DMRs

In neuronal cells and human tissues, some CG-DMRs were found in large clusters, forming hundreds of kilobase-scale hypomethylated domains, termed “large hypo CG-DMRs”^9,44^. To identify this methylation feature in fetal tissues, for each tissue type, we merged tissue-specific CG-DMRs that are within 1kb and filtered for ones larger than 2kb, as previous described^44^. Consistent with previous results^44^, the large hypo CG-DMRs (n=273-1,302), which comprise of 0.7%-1.6% the merged CG-DMRs, showed higher levels of H3K4me1 and H3K27ac compared with the typical smaller CG-DMRs, implying that these clusters of predicted fetal enhancers may represent tissue-specific super-enhancers^54^ (Fig. 4a-e). Indeed, 25-57% of the large hypo CG-DMRs overlapped with regions predicted as super-enhancers (Fig. 4f; Methods). Like super-enhancers, we found that a very large portion (58%-79%) of these large hypo CG-DMRs are intragenic (p-value < 0.001; Methods) and are significantly associated with genes enriched for tissue-related biological processes (false discovery rate < 0.01; Supplemental Table 8).

**Figure 4.**
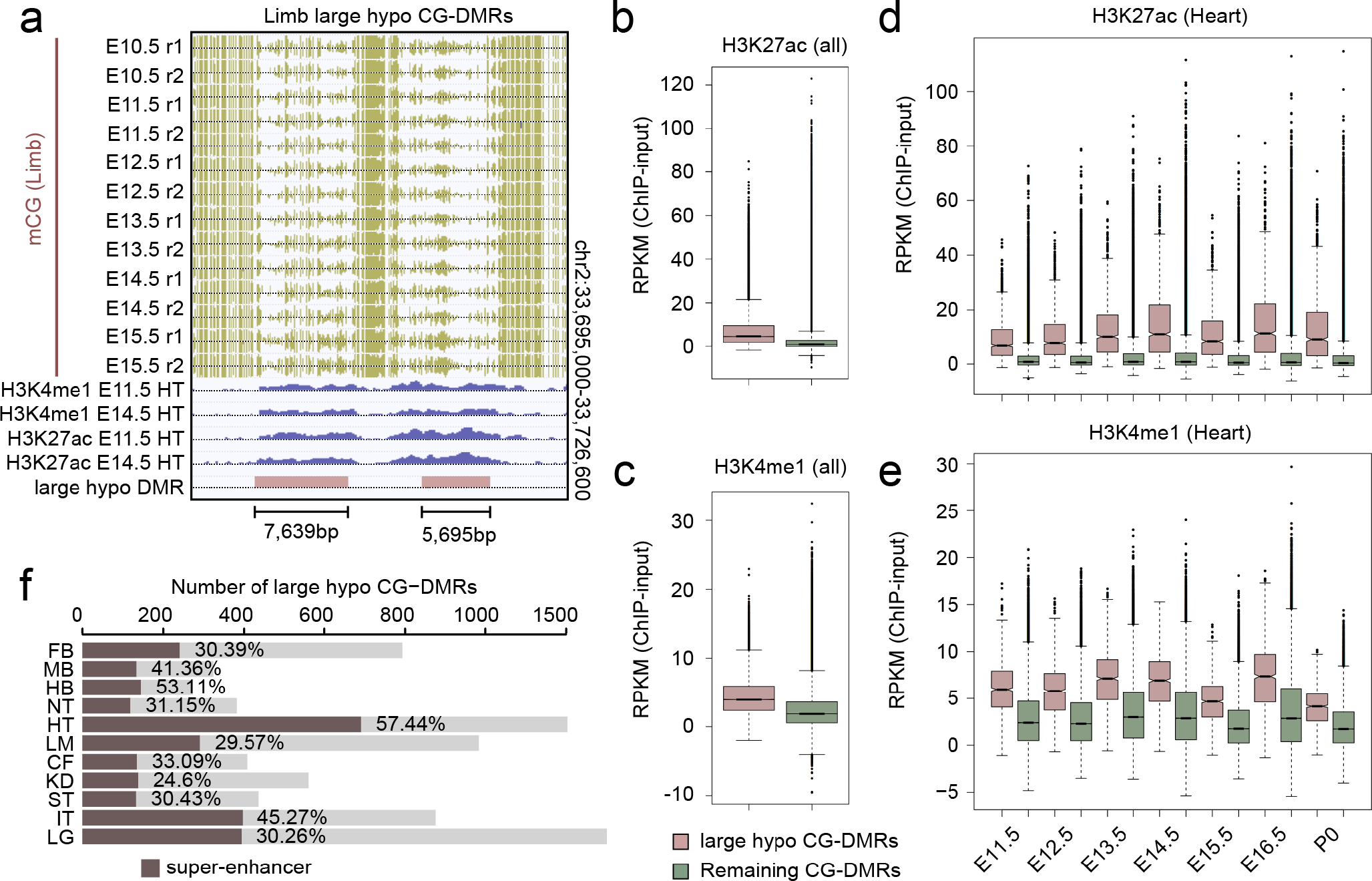
Large hypo CG-DMRs are enriched for super-enhancers. **a**, Examples of large hypo CG-DMRs. **b-c**, H3K27ac (b) and H3K4me1 (c) enrichment in large hypo CG-DMRs (red) and the rest of CG-DMRs (green) at all developmental stages across all tissue types except liver. **d-e**, H3K27ac (d) and H3K4me1 (e) enrichment in heart-specific large hypo CG-DMRs (red) and the rest of heart-specific CG-DMRs (green) in heart tissue from all developmental stages. **f**, Number of large hypo CG-DMRs identified in each tissue type and the percentage that overlap with super-enhancers (red).

These large hypo CG-DMR are different from another type of multi-kilobase DNA methylation feature called the DNA methylation valley (DMV; Methods)^55^. DMVs are generally ubiquitously lowly methylated in all tissues across developmental trajectory (Extended Data Fig. 8a-b). Large hypo CG-DMRs, in contrast, display striking spatiotemporal hypomethylation patterns (Extended Data Fig. 8a, c). Consistently, less than 4% of large hypo CG-DMRs overlapped with DMVs. Furthermore, we found that 53-58% of the DMV-overlapped genes encoding TFs, much more than large hypo CG-DMRs (8-17%; Extended Data Fig. 8d).

### Intragenic non-CG methylation

A less well-understood form of cytosine DNA methylation found in mammal genomes is called non-CG methylation or mCH^18^. While mCH is only present at low levels in most adult tissues, it is the dominant form of cytosine methylation in human neurons^18^. While no mechanism is known to actively remove mCH, passive dilution can occur during cell replication^18^. Surprisingly, we found that mCH accumulates to detectable levels in nearly all fetal tissues as some point during their developmental trajectories; the timing varies in different tissues/organs (Fig. 5a). For example, three brain distinct region tissues showed different rates of mCH accumulation the timing of which correlates with their developmental maturation (lower expression of neural progenitor markers and higher expression of neuronal markers) in sequential order of hindbrain, midbrain, and forebrain (Fig. 5a; Extended Data Fig. 9). mCH is preferentially deposited at 5’-CAG-3’ context in embryonic stem cells and at 5’-CAC-3’ context in adult tissues^8–10,17–19^. In all fetal tissues we found mCH enriched in the CAC context and its enrichment in this 3 base specificity further increased as the tissues mature, implying a similar DNMT3A-dependent mCH pathway in both fetal and adult tissues^18^ (Extended Data Fig. 10a).

**Figure 5.**
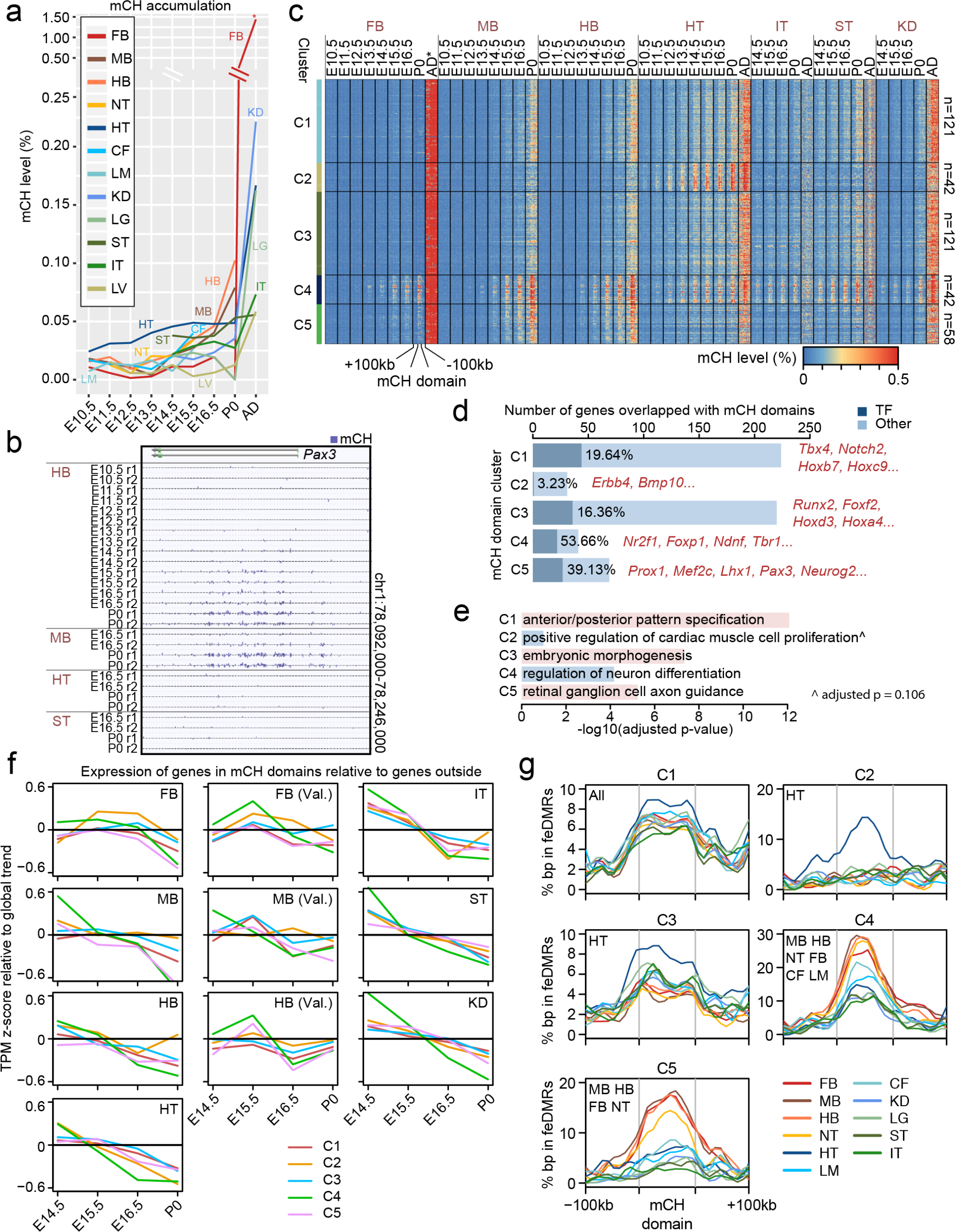
mCH accumulation predicts reduced gene expression. **a**, Genome-wide non-CG methylation (mCH) levels for each tissue across their developmental trajectories. The adult (AD) forebrain data (postnatal 6 week frontal cortex) is from Lister et al^10^. **b**, An example of a mCH domain. Enriched for mCH accumulation determined by comparison to flanking regions. **c**, K-means clustering identification of 384 mCH domains clustered into 5 groups based on the tissue-specific mCH accumulation. Heatmap showing the methylation profiling of mCH domains and flanking genomic regions (100kb upstream and 100kb downstream). **d**, Number of genes overlapping mCH domains in each of 5 groups. Dark blue bars indicate the number of genes encode transcription factors in mCH domains. Examples of genes located within mCH domains are listed on the right. **e**, The most enriched gene ontology (Biological Process) terms for genes that lie within mCH domains for each cluster. **f**, Expression dynamics of genes within mCH domains relative to the other genes. Z-scores were calculated for each gene across development and each line shows the mean value of mCH overlapping genes for each cluster. **g**, Tissue-specific enrichment of feDMRs in mCH domains. Each mCH domain was divided into 10 bins and its flanking regions included ten 10kb upstream bins and ten 10kb downstream bins. Line plots show the fraction of bases in each bin that are overlapped with tissue-specific feDMRs. A list of tissues in each plot indicates wthere feDMRs are enriched in mCH domains compared to flanking regions (from the most enriched to the least enriched).

Intriguingly, mCH was not uniformly distributed across the genome of fetal tissues but preferentially accumulated at genomic regions that we termed as “mCH domains”; genomic locations showing higher mCH levels than their flanking sequences (Fig. 5b). In total, 384 mCH domains were identified, which averaged 255kb in length (Methods). Strikingly, 92% (355 out of 384) of mCH domains and 61% of the bases within them overlapped with annotated gene bodies (p-value < 0.001; Methods). Interestingly we found a highly significant fraction (22%) of the mCH domain genes (e.g. *Pax3*) encode transcription factors, many of which are known to be involved in tissue development/organogenesis (Fig. 5b; 128 out of the 581 genes, p-value < 0.001; Methods).

To further explore the tissue and temporal specificity of mCH accumulation, we used k-means clustering to group mCH domains into 5 clusters based on methylation dynamics (Fig. 5b-c; Extended Data Fig. 10b; Methods). mCH domains in clusters 1 and 3 acquire mCH in all tissues and are enriched for genes related to embryo development (Fig. 5c-e; Extended Data Fig. 10b; Supplemental Table 9). mCH accumulation in cluster 4 is also observed in all tissues but is more pronounced than in clusters 1 and 3. Interestingly, the cluster 4 mCH domains were significantly enriched for genes involved with neuron differentiation functions (Fig. 5d-e). In contrast to these ubiquitous mCH domains, tissue-specific mCH domains were also found. Cluster 2 gains mCH mostly in heart where as those in cluster 5 show brain-specific mCH accumulation, overlapping genes that are enriched for functions related to axon guidance (Supplemental Table 9).

As mCH accumulates in mCH domains at late developmental stages, the genes within tend to be repressed in their expression compared with genes outside mCH domains, especially by P0 (Fig. 5f; Extended Data Fig. 10c). Since mCH domains are enriched for TFs and other genes related to tissue/organ or embryo development, our data suggests that mCH may be associated with silencing of the pathways of early fetal development.

Interestingly, mCH domains showed enrichment for tissue-specific feDMRs compared to flanking regions (Fig. 5g). feDMRs in mCH domains tend to lose enhancer-like chromatin signatures as development proceeded (decreasing enhancer score; Extended Data Fig. 10d). In addition, these feDMRs, in late stages, tend to become hypermethylated compare with ones outside mCH domains (Extended Data Fig. 10e).

These observations indicate that mCH accumulation predicts the future silencing of regulatory elements, consistent with recently reported findings for human cerebral organoids^56^. Collectively, we found mutual associations between several mCH domains features including: mCH accumulation, enrichment for genes related to the tissues that acquire mCH, down-regulation of gene expression and enrichment of feDMRs. Delineating the mechanism(s) that drive these associations will provide new insights into mCH regulation and the potential involvement of non-CG methylation in transcriptional regulation.

## Conclusion

In this study we describe the generation and analysis of a comprehensive collection of base-resolution, genome-wide maps of cytosine methylation for 12 fetal tissue types from 8 distinct developmental stages of mouse embryogenesis. By integrating DNA methylation, histone modification and RNA sequencing data from the same tissue samples, we annotated millions of methylation variable elements including prediction of fetal enhancers and sets of transcription factors that bind them. The human counterparts of these fetal enhancers are tissue-specifically enriched for genetic risk loci associated with human diseases. Such enrichments suggest the possibility of generating mouse models of human diseases by introducing the specific alleles into feDMRs using genomic editing techniques^57^. Because of the temporal nature of these data, we uncovered surprisingly simple mCG dynamics at predicted DNA regulatory regions. During early stages of fetal development, methylation decreases at predicted fetal regulatory elements in all tissues until birth, after which time it dramatically rises. Since the tissues that we have investigated are composed by a variety of cell types, a fraction of the observed dynamics potentially result from DNA methylation changes during the differentiation of individual cell types and/or the changing cell type composition during development. In spite of the tissue heterogeneity, such dynamics suggest a plausible regulatory principal whereby stable repressive mCG is removed to enable more rapid, flexible modes of gene regulation (e.g. histone modification/open chromatin). Also, our findings extend current knowledge of methylation in a non-CG context, an understudied type of DNA methylation. We observed that during fetal development there is preferential accumulation of mCH tissue-specifically at genomic locations, each hundreds of kilobases in size, a novel genomic feature we termed “mCH domains”. Genes that lie in mCH domains become down-regulated in their expression as mCH further accumulates during the later stages of fetal development. Though its function remains debatable, *in vivo* and *in vitro* studies indicated that mCH directly increases the binding affinity of MeCP2^21–23^, mutation of which leads to Rett Syndrome. Gene-rich mCH domains are likely enriched for yet to be discovered mCH binding proteins, which, like MeCP2, may be involved recruiting transcriptional repressor complexes, promoting the observed gene repression. Our study highlights the power of temporal tissue epigenome maps to uncover regulatory element dynamics in fetal tissues during *in utero* development. These spatiotemporal epigenomic datasets provide a valuable resource for studies of fundamental questions about gene regulation during mammalian tissue/organ development as well as knowledge about the possible origins of human developmental diseases.

## Acknowledgements

We thank Drs. Junhao Li, Shao-shan Carol Huang, Eran A. Mukamel and Liang Song for critical comments. Y.H. is supported by the H.A. and Mary K. Chapman Charitable Trust. D.U.G is supported by the A.P. Giannini Foundation and NIH IRACDA K12 GM068524. A.V. and L.A.P. was supported by National Institutes of Health grants R01HG003988, U54HG006997, R24HL123879 and UM1HL098166, and the research was conducted at the E.O. Lawrence Berkeley National Laboratory and performed under Department of Energy Contract DE-AC02-05CH11231, University of California. J.R.E is an Investigator of the Howard Hughes Medical Institute. This work used the Extreme Science and Engineering Discovery Environment (XSEDE), which is supported by National Science Foundation grant number ACI-1548562. This work was supported by the National Institutes of Health ENCODE Project (U54 HG006997). The data that support the findings of this study are publically accessible at https://www.encodeproject.org/search/?searchTerm=ecker&type=Experiment&award.rfa=ENCODE3 and http://neomorph.salk.edu/ENCODE_mouse_fetal_development/. The additional RNA-seq dataset of forebrain, midbrain, hindbrain and liver is available at the NCBI Gene Expression Omnibus (GEO) under accession GSE100685. Details of data used in this study can be found in Supplemental Table 1 and 2.

## Contributions

Y.H., M.H., R.F., H.C., B.L. performed data analysis. Y.H. and M.H. wrote the manuscript. C.L. and J.R.E edited the manuscript. D.E.D., A.V. and L.A.P. group collected the tissues from E11.5 mouse embryo, which were later under epigenome and transcriptome profiling. D.U.G, A.Y.L, and B.R. generated the histone modification data. J.R.N. and R.C generated the whole genome bisulfite sequencing data and the validation set of RNA-seq data. B.A.W., D.T. and H.A. generated the RNA-seq data. J.R.E supervised the project.

## Supplemental Note - mCG landscape remodeling in fetal liver

Distinct from the hypermethylated genome of all other tissues, the liver genome underwent drastic global demethylation from E11.5 to E14.5, and remained hypomethylated till E16.5, after which it returned to hypermethylated state at P0 (Fig. 1B). The hypomethylated liver genome, present during E12.5 to E16.5, displayed a partially methylated domains (PMDs) signature, a methylation feature previously observed in human cultured cell lines^8,29^, pancreas^9^, placenta^30^ and cancer samples^31,32^. PMDs are large genomic regions (typically greater than 100kb) that are lowly CG methylated (Extended Data Fig. 1A). We systematically identified PMDs in liver samples from all stages (Methods). Strikingly, from E14.5 to E16.5, PMDs covered more than half of the genome and the coverage shrunk dramatically afterwards (Extended Data Fig. 1B). We found that PMDs identified in E15.5 displayed hypomethylation in all liver samples and covered almost all PMDs from other stages (Extended Data Fig. 1C-D). These results indicate that the PMDs identified at different fetal stages are essentially identical and the different PMDs calls were due to various signal-to-noise ratios. Therefore, we defined the PMDs identified in E15.5 liver as liver PMDs (n = 4,578; average size = 338kb).

Mouse liver PMDs share all molecular signatures of the PMDs identified in human fibroblast cell lines, normal and cancer tissues^8,9,31,32^: First, mouse PMDs are enriched for H3K9me3 and H3K27me3 and are depleted of H3K27ac (Extended Data Fig. 1E). Second, mouse PMDs tend to be replicated during the later stages of the cell cycle and strongly overlap with lamina-associated domains (Extended Data Fig. 1F; p-value < 0.001; permutation test; Methods). Furthermore, we found that genes overlapping with mouse PMDs tend to have lower expression compared to genes outside PMDs, which was also reported by Schultz et al for human pancreas^9^. These shared properties indicate that human and mouse PMDs are likely identical genome feature and their presence may be due to similar mechanism, likely the failure of mCG maintenance in rapidly dividing cells^31^.

The presence of PMDs in fetal liver coincides with hematopoiesis^61,62^. Hematopoiesis initiates at E11.5, while liver genome remains hypermethylated (Fig. 1B). Then, hematopoietic expansion occurs between E12.5 and E14.5, during which the liver genome underwent demethylation and PMDs became evident (Fig. 1B; Extended Data Fig. 1B and D). The increasing number of rapidly dividing cells during this expansion period may explain the formation of PMDs. After E15.5, the hematopoiesis starts to disappear although the “liver tissue” genome is not fully remethylated until P0 (Extended Data Fig. 1B).

**Extended Data Figure 1.**
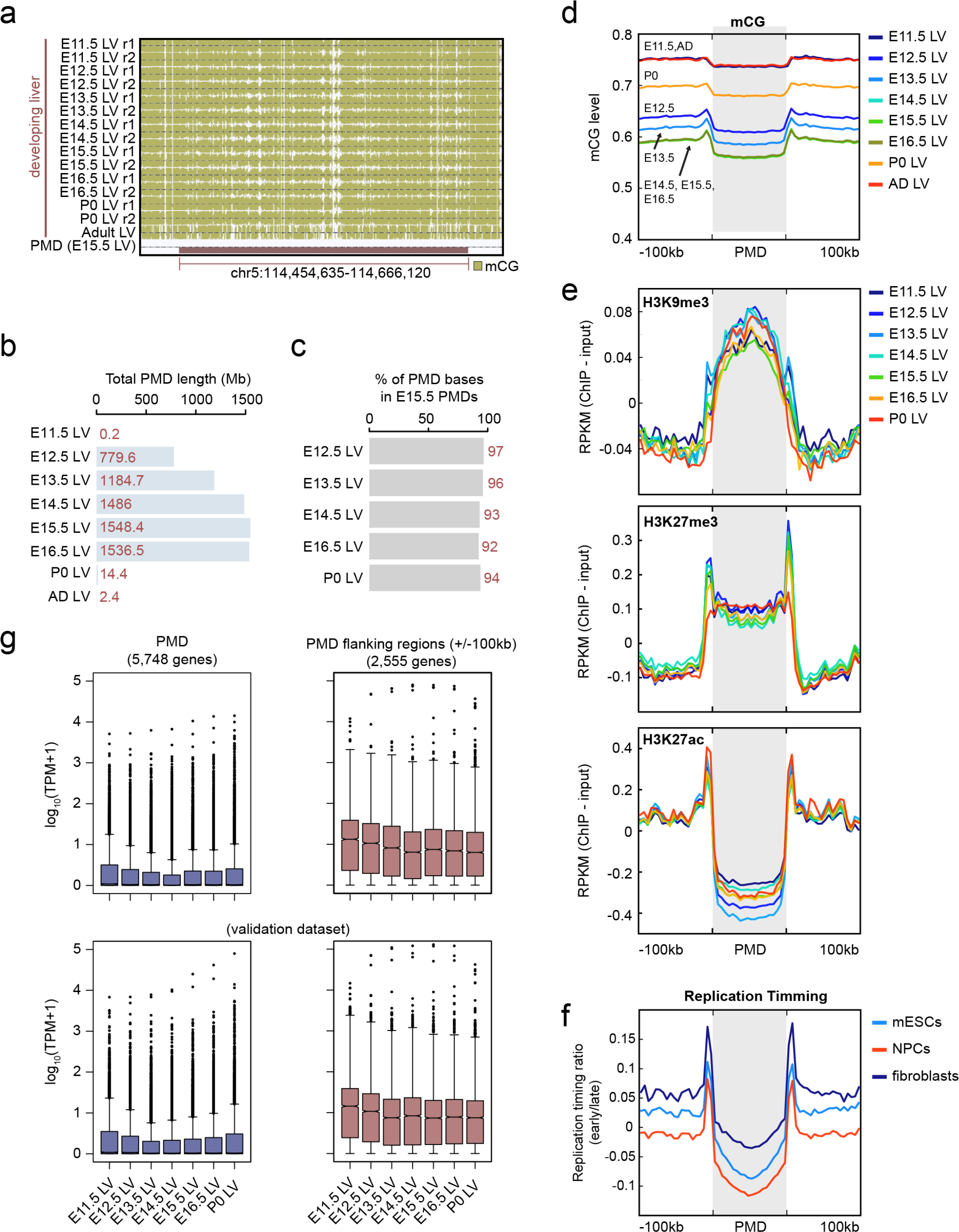
Global hypomethylation in fetal liver. **a**, Example of a partially methylated domain (PMDs) in developing mouse fetal liver. The PMD location is marked by a red bar. **b**, The total bases that PMDs encompass in liver at different developmental stages. **c**, Percentage of bases in the PMDs identified in each of the liver samples (E12.5 liver, E13.5 liver etc) that are also within the PMDs identified in E15.5 liver sample. **d**, Average mCG level (mCG/CG) of PMDs and flanking regions (+/-100kb) in liver samples from different developmental stages. **e**, Histone modification profiles for H3K9me3 (top), H3K27me3 (middle) and H3K27ac (bottom) within PMDs and flanking regions (+/-100kb) in liver samples from different developmental stages. **f**, Replication timing profiling of PMDs and flanking regions (+/-100kb). The values indicate the tendency to be replicated at an earlier stage in the cell cycle. **g**, Expression of genes overlapping PMDs and flanking regions (+/-100kb) (left) compared with those with no PMD overlap (right). Two plots on the bottom show the data from a validation dataset, containing RNA-seq data generated using different protocol on the matched tissues.

**Extended Data Figure 2.**
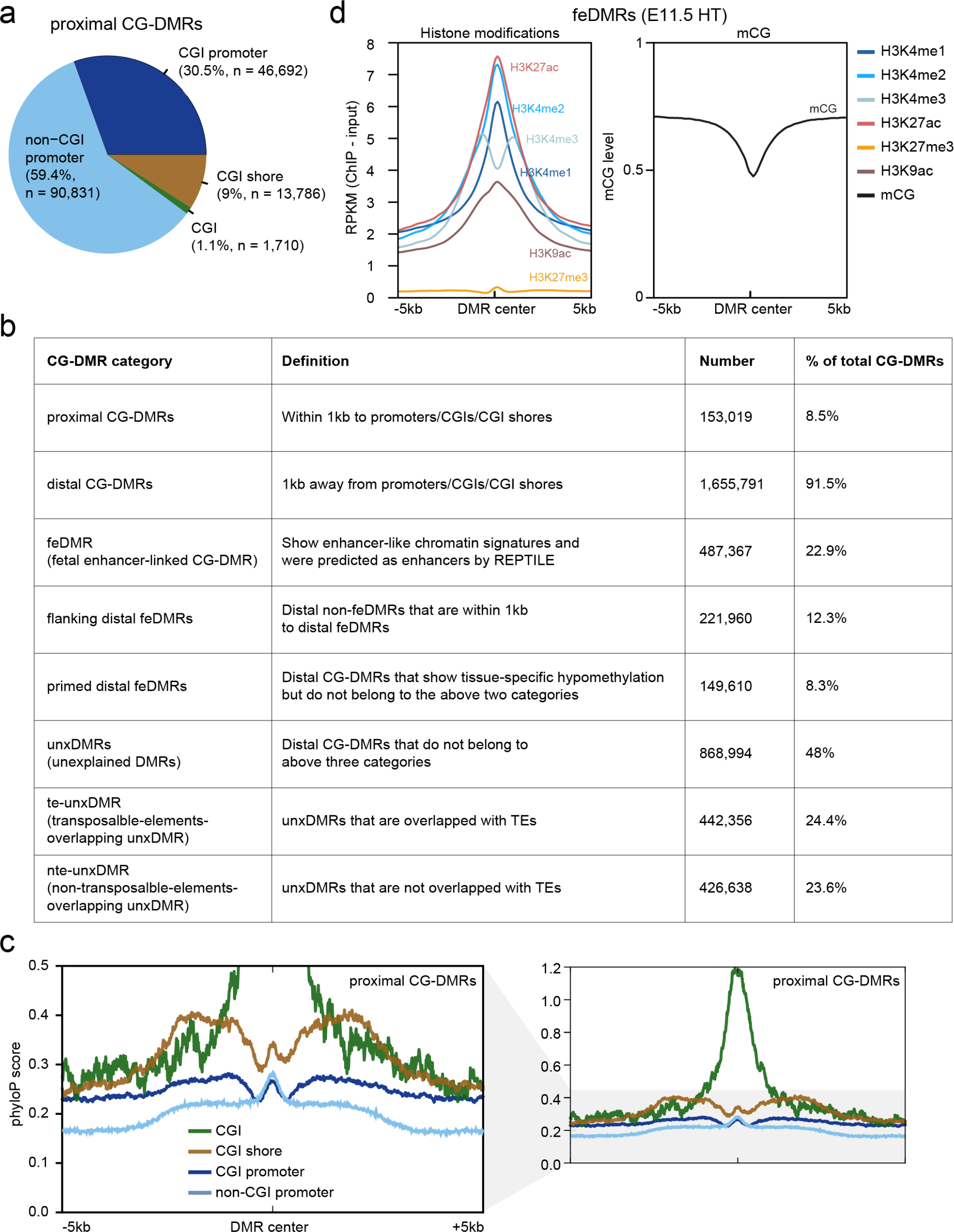
Categorization of CG-DMRs. **a**, Genomic distribution of proximal CG-DMRs. **b**, Evolutionary conservation of proximal CG-DMRs overlapping with: CG islands (CGI), CGI shores, CGI promoters and non-CGI promoters. phyloP score was used to measure the degree of conservation. **c**, Chromatin signatures of fetal enhancer-linked CG-DMRs in E11.5 heart. The aggregate line plots show the average histone modification and mCG profiles of +/-5kb regions centered at CG-DMR centers. **d**, Table summarizing the definition and number of various CG-DMR categories. Note that categories in the table are mutually exclusive.

**Extended Data Figure 3.**
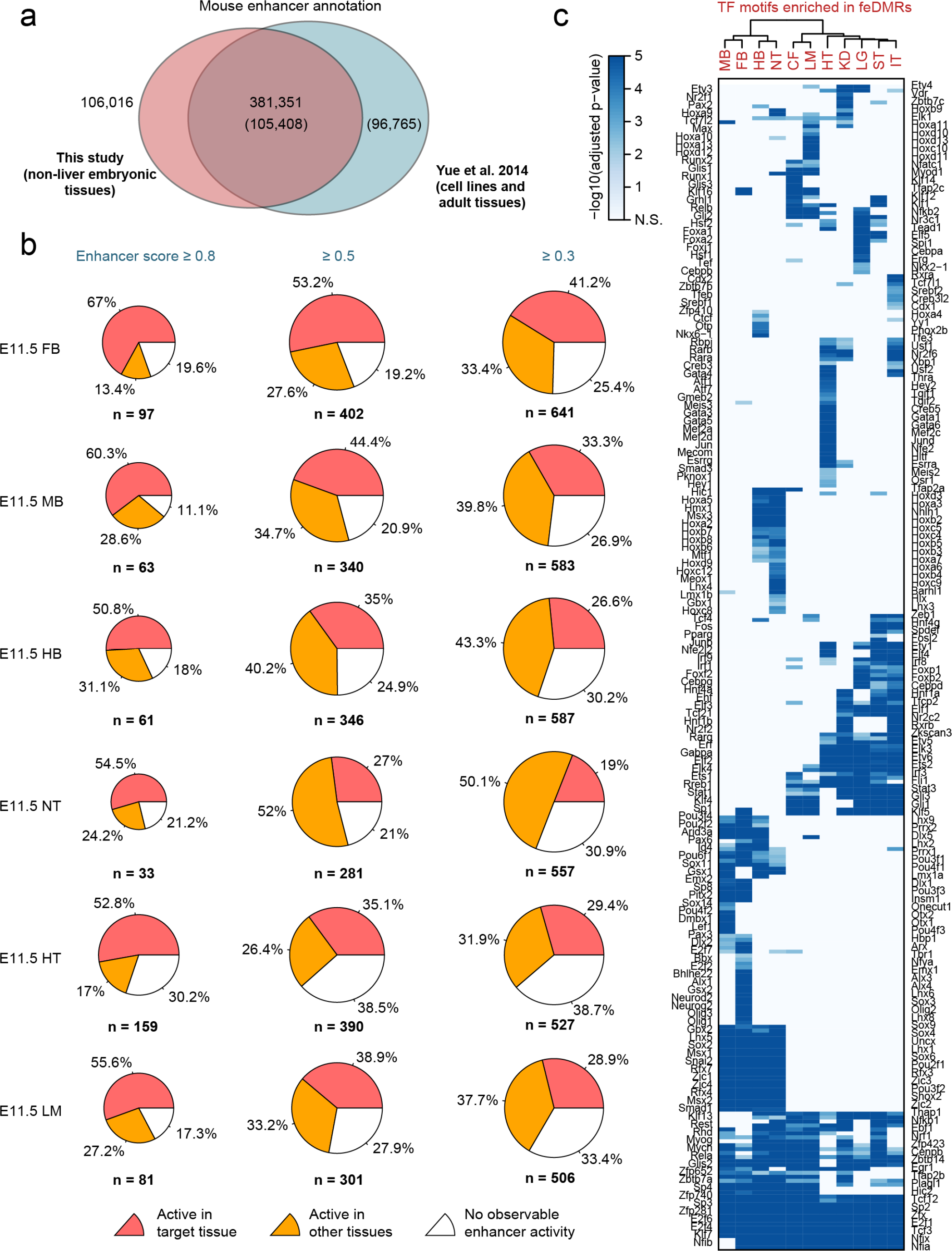
Fetal-enhancer-linked CG-DMRs (feDMRs) **a**, The overlap between feDMRs identified in this study and the enhancers predicted in Yue et al^37^. Numbers in parenthesis indicate the counts of enhancers from Yue et al, whereas the remaining numbers denote the counts of feDMRs. **b**, Experimental validation results of the feDMR-overlapping elements from VISTA enhancer browser^38^. Different enhancer score thresholds were used for calling feDMRs for each tissue at fetal stage E11.5. Each pie shows the fraction of elements that were experimentally validated as active enhancers in matched tissue (red) or any tissue (orange) or as inactive enhancers (white) at fetal state E11.5. **c**, Enrichment of transcription factor binding motifs in feDMRs of different tissue types.

**Extended Data Figure 4.**
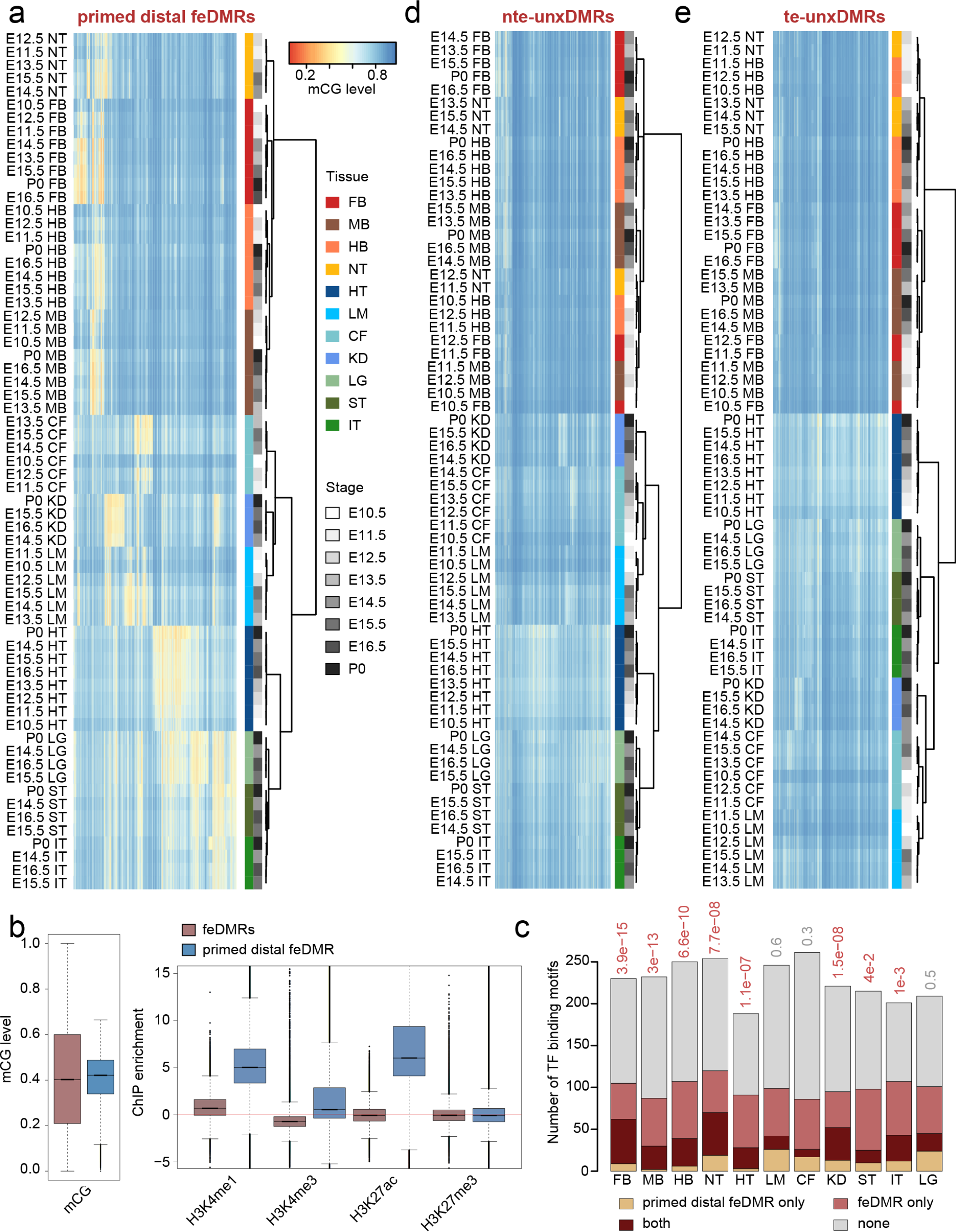
Characterization of primed distal feDMRs and unxDMRs. **a**, CG methylation (mCG) level of all primed distal fetal enhancer-linked CG-DMRs (feDMRs) in all non-liver tissues. Each row in the heatmap is one tissue sample and each column corresponds to one primed distal feDMR. Both rows and columns were clustered using hierarchical clustering. Colors bars indicate the tissue types and developmental stages of samples, respectively. **b**, mCG (left) and histone modification (right) signatures of primed distal feDMRs (blue) and feDMRs (red). Boxplots show the median and quantiles of the values in all non-liver tissues. **c**, Number of enriched transcription factor binding motifs only in feDMRs (red), only in primed distal feDMRs (orange), both (dark red) and none (grey). Only the motifs linked to expressed transcription factors (transcripts per million, TPM >= 10) were included. **d-e**, Similar to (a), heatmaps showing the mCG levels of unexplained CG-DMRs, including te-unxDMRs (d) and nte-unxDMRs (e).

**Extended Data Figure 5.**
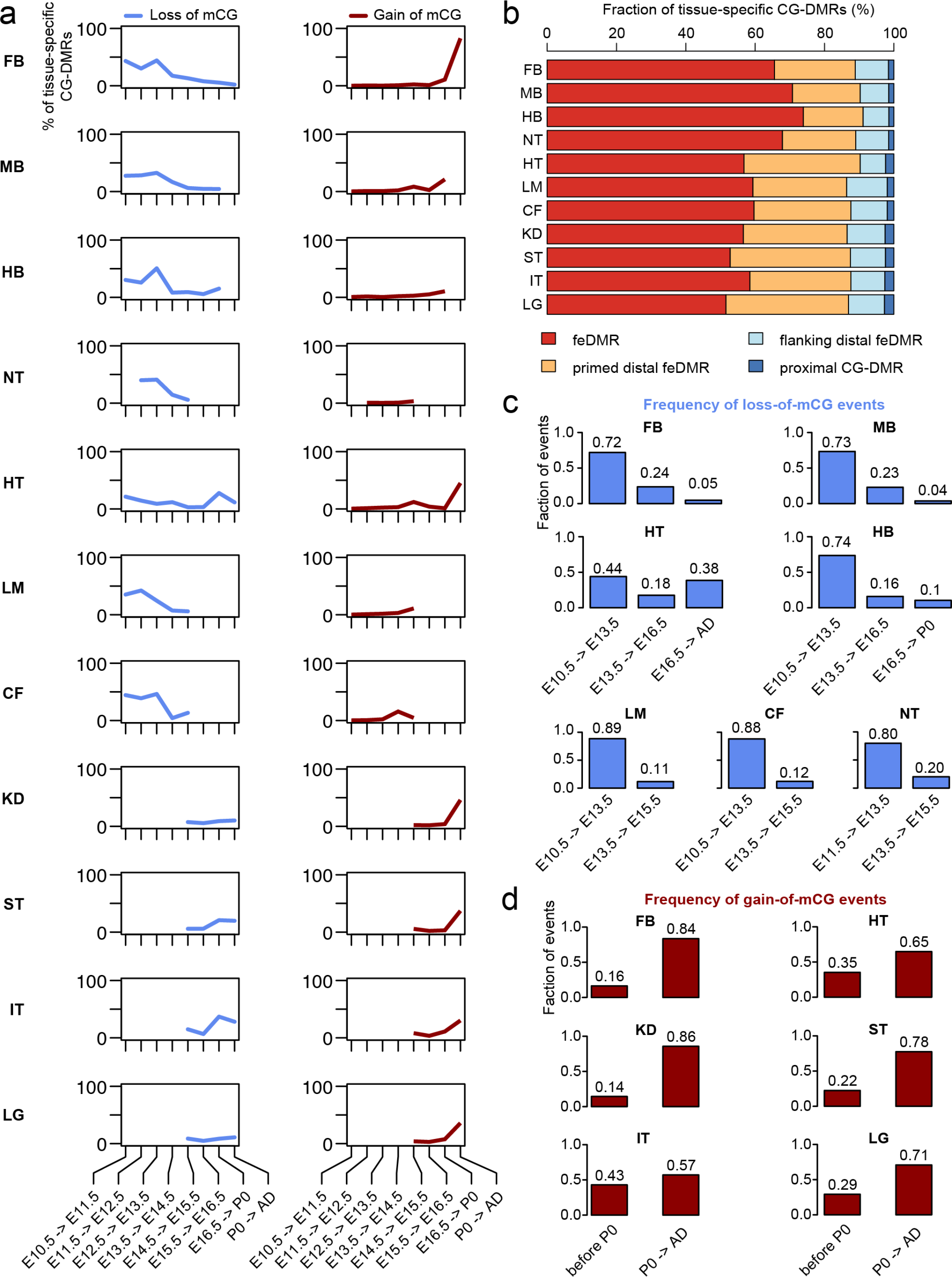
mCG dynamics of tissue-specific CG-DMRs. **a**, Fraction of tissue-specific CG-DMRs showing loss-of-mCG (blue) or gain-of-mCG (red) for each fetal stage. Loss-of-mCG (gain-of-mCG) event is defined as an increase (decrease) of mCG of at least 0.1. **b**, Composition of tissue-specific CG-DMRs. **c**, Percentage of loss-of-mCG events for each fetal stage. **d**, Percentage of gain-of-mCG events for each fetal stage.

**Extended Data Figure 6.**
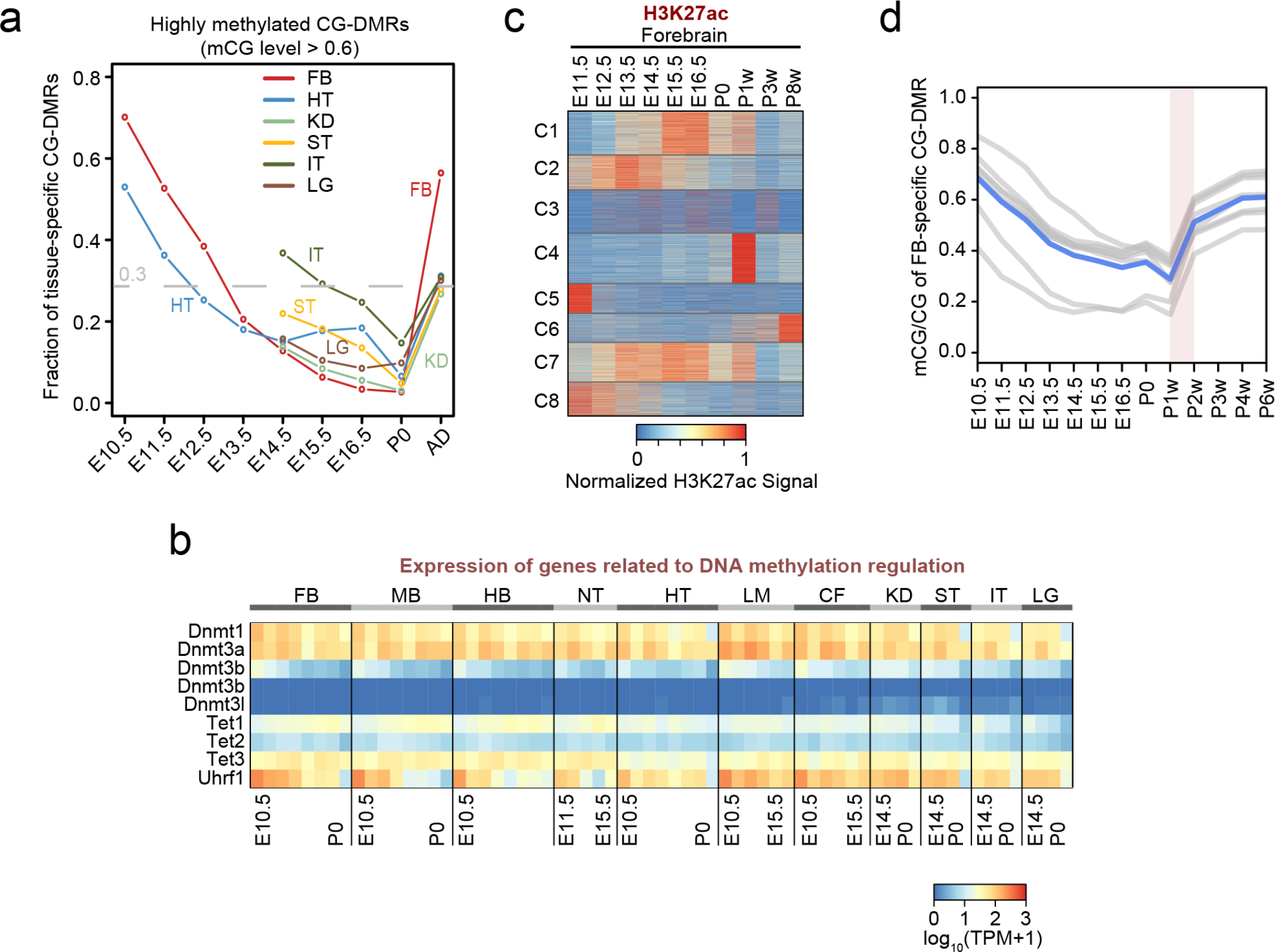
Link between methylation dynamics and histone modifications at tissue-specific CG-DMRs. **a**, Fraction of tissue-specific CG-DMRs that are heavily CG methylated (mCG level > 0.6). **b**, RNA abundance of genes involved in DNA methylation pathways, measured by transcripts per million (TPM). **c**, Normalized H3K27ac signals in different clusters. **d**, Dynamic mCG level of forebrain-specific CG-DMRs. Grey lines show the mean methylation levels of CG-DMRs in different clusters. Blue line is the mean of all clusters (grey lines).

**Extended Data Figure 7.**
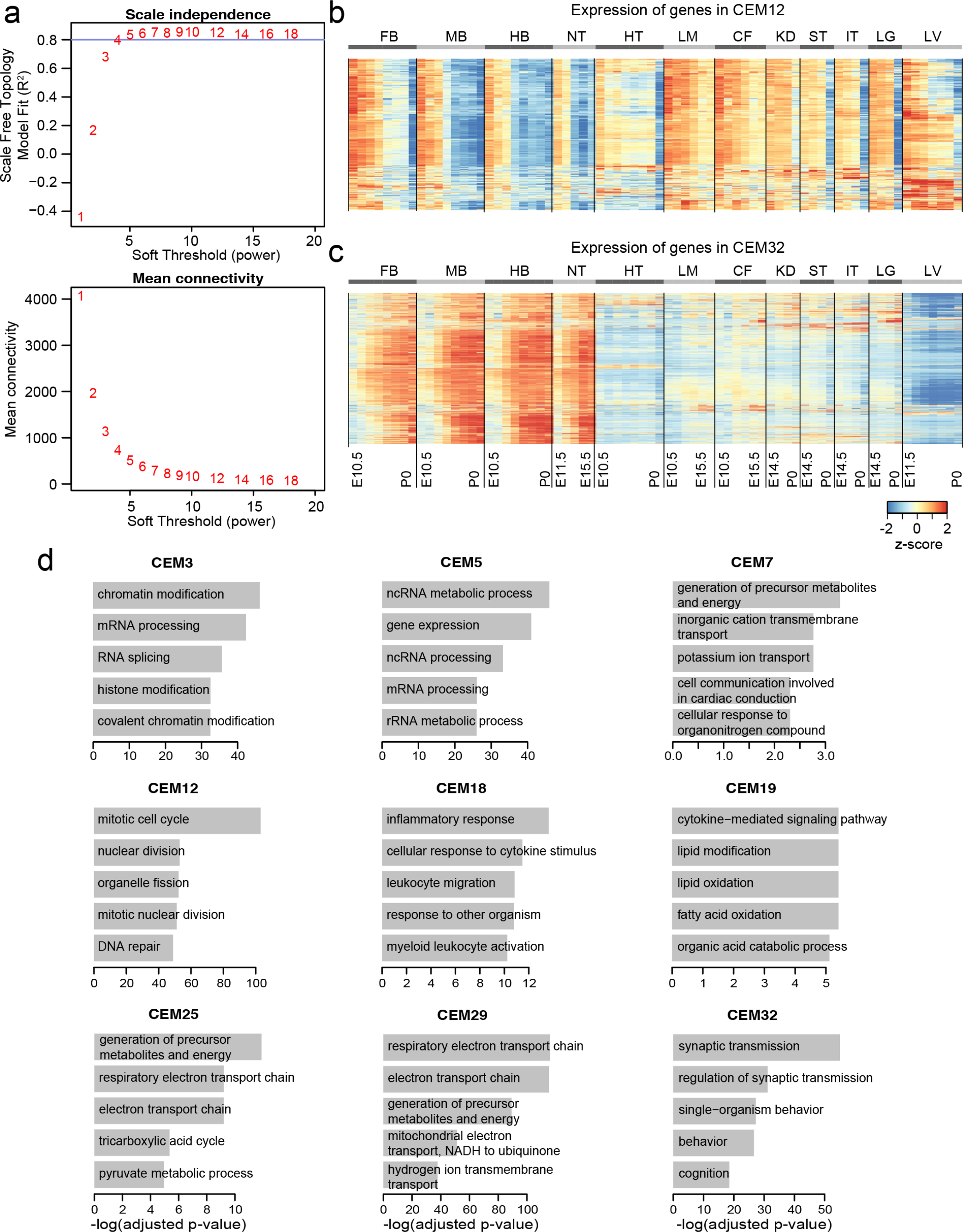
WGCNA identification of co-expression modules. **a**, The scale free topology model fit (R^2^) (top) and the mean connectivity of the coexpression network (bottom) given different soft-thresholding powers. These two plots show how thresholds were chosen for weighted gene co-expression network analysis (WGCNA). Blue horizontal line indicates the model fit cutoff (R^2^ = 0.8). A soft threshold = 5 was chosen to construct the co-expression network because it is first threshold value where the model fit is greater than 0.8. **b-c**, Expression of genes in CEM12 (b) and CEM32 (c). Each row is a gene in certain module and the transcripts per million (TPM) z-scores were calculated along each row. d, Top enriched ontology terms of genes in co-expression modules.

**Extended Data Figure 8.**
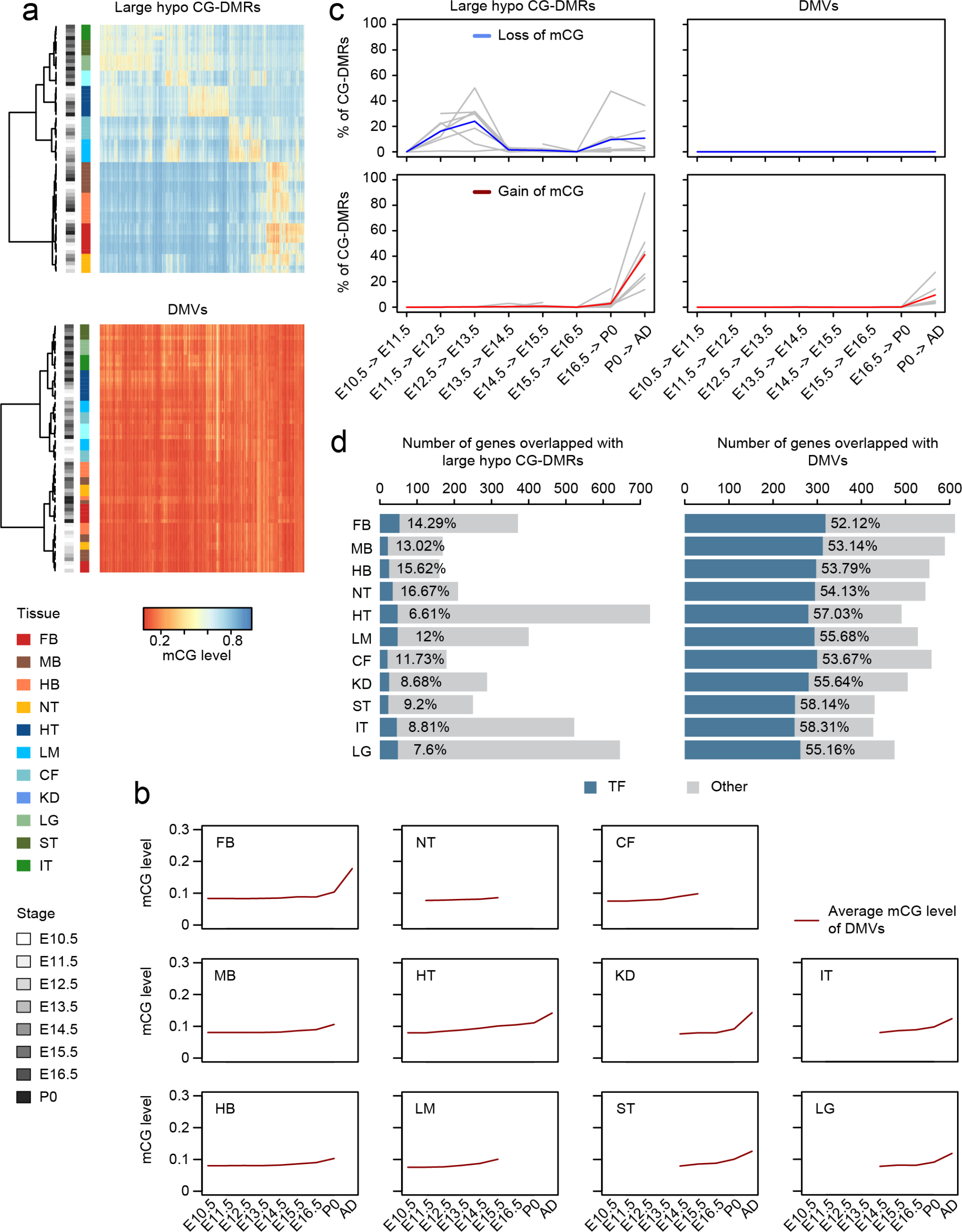
Comparing large hypo CG-DMRs and DNA methylation valleys (DMVs). **a**, CG methylation (mCG) level of large hypo CG-DMRs (top) or DMVs (bottom) in all non-liver tissues. Both rows and columns were clustered using hierarchical clustering. Colored bars indicate the tissue types and developmental stages of samples, respectively. Heatmap shows the data of merged large hypo CG-DMRs (DMVs) for predictions from all tissue samples. See Methods for details. **b**, Fraction of large hypo CG-DMRs (left) or DMVs (right) that undergo lost-of-mCG (top, blue) and gain-of-mCG (bottom, red) during development. The blue (loss-of-mCG) or red (gain-of-mCG) line shows the aggregated values over all non-liver tissues, whereas grey lines show the data for each tissue type. **c**, Average mCG level of DMVs identified in each tissue type. **d**, Number of genes overlapping with large hypo CG-DMRs (left) or DMVs (right). Dark blue indicates genes encoding transcription factors (TFs).

**Extended Data Figure 9.**
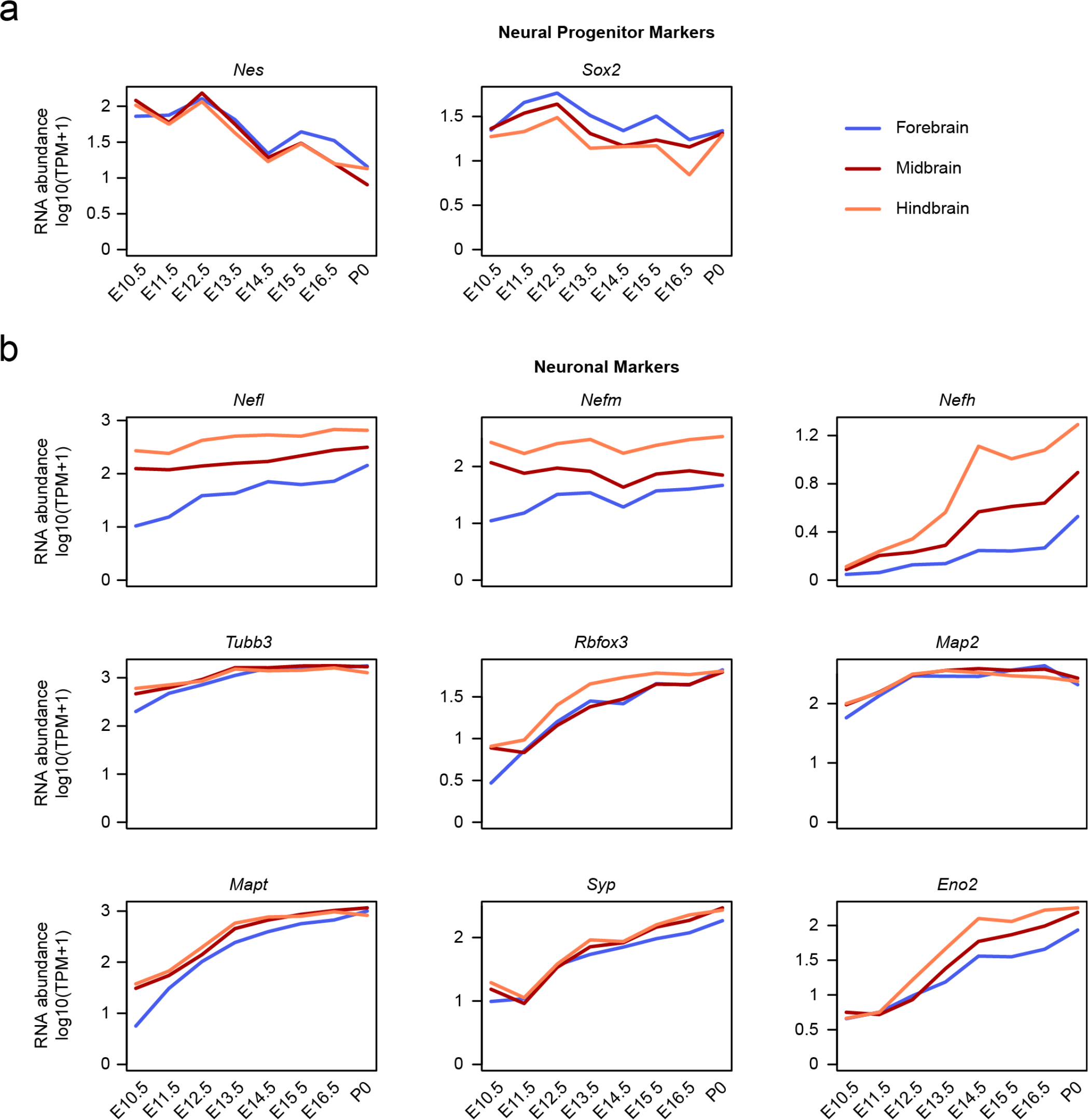
Expression of neural progenitor and neuronal markers in different brain regions. **a**, Expression of neural progenitor markers, *Nes* ^58^ and *Sox2*^59^. TPM – transcripts per million mapped reads. **b**, Expression of several neuronal markers from Svendsen et al^60^.

**Extended Data Figure 10.**
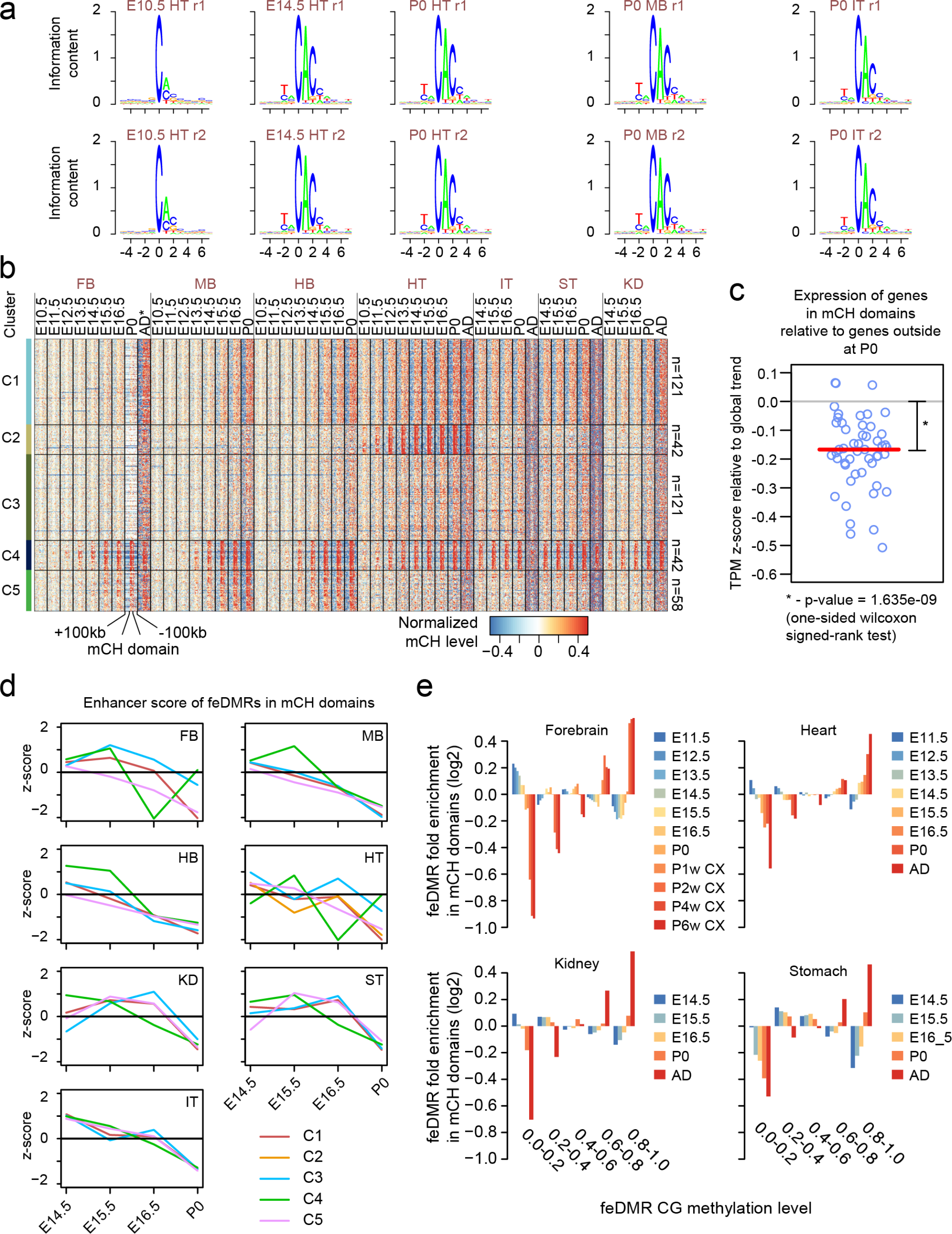
Non-CG methylation accumulation in fetal tissues. **a**, Sequence context preference for non-CG methylation (mCH). **b**, Grouping mCH domains into 5 clusters based on the dynamics of methylation accumulation. The heatmap shows normalized methylation levels of mCH domains and flanking genomic regions (up to 100kb upstream and 100kb downstream). mCH in the adult (AD) forebrain was approximated using data of frontal cortex from 6-week-old mice. **c**, Average enhancer score dynamics of feDMRs within mCH domains. Z-scores were calculated for each feDMR across development and each line shows the mean value of the mCH domains overlapping feDMRs for each cluster. **d**, Enrichment of tissue-specific feDMRs that showed different CG methylation levels in mCH domains. Colors (from blue to red) denote the stages where the mCG level was calculated. **f**, The expression of genes in mCH domains at P0 relative to the expression dynamics of genes outside mCH domains. Each circle corresponds to the value given one mCH domain cluster and one tissue. Red horizontal line indicates the median, which was tested against 0 using one-sided Wilcoxon signed-rank test.

## Methods

### Data Availability

All whole-genome bisulfite sequencing (WGBS) data from mouse embryonic tissues are available at the ENCODE portal (https://www.encodeproject.org/) and also deposited in the NCBI Gene Expression Omnibus (GEO) (Supplemental Table 1). The additional RNA-seq dataset of forebrain, midbrain, hindbrain and liver is available at the NCBI Gene Expression Omnibus (GEO) under accession GSE100685. All other data used in this study, including chromatin immunoprecipitation sequencing (ChIP-seq), RNA-seq and additional WGBS data, are available at the ENCODE portal and/or GEO (Supplemental Table 2).

### Code availability

Custom code used for this study is available upon request. This work used computation resource from the Extreme Science and Engineering Discovery Environment (XSEDE)^63^.

### Abbreviations

- AD: adult
- CEM: co-expression module
- mC: cytosine DNA methylation
- mCG: CG methylation
- mCH: non-CG methylation
- TF: transcription factor
- H3K4me1: Histone 3 lysine 4 monomethylation
- H3K4me3: Histone 3 lysine 4 trimethylation)
- H3K27me3: Histone 3 lysine 27 trimethylation
- H3K27ac: Histone 3 lysine 27 acetylation
- WGBS: whole-genome bisulfite sequencing
- REPTILE: Regulatory Element Prediction based on TIssue-specific Local Epigenetic marks
- GWAS: genome-wide association study
- SNP: single nucleotide polymorphism
- TPM: transcripts per million
- WGCNA: weighted gene co-expression network analysis
- RPKM: Reads Per Kilobase per Million mapped reads

#### Genomic features

- CG-DMR: differentially CG methylation region
- feDMR: fetal enhancer linked CG-DMR
- fd-feDMR: flanking distal feDMR
- pd-feDMR: poised distal feDMR
- unxDMR: unexplained CG-DMR
- te-unxDMR: transposable element overlapping unxDMR
- nte-unxDMR: non transposable element overlapping unxDMR
- TSS: transcription start sites
- CGI: CpG island
- DMV: DNA methylation valley
- PMD: partially methylated domain
- AD-A enhancer: adult active enhancer
- AD-V enhancer: adult vestigial enhancer

#### Tissues/organs

- FB: forebrain
- MB: midbrain
- HB: hindbrain
- NT: neural tube
- HT: heart
- CF: craniofacial
- LM: limb
- KD: kidney
- LG: lung
- ST: stomach
- IT: intestine
- LV: liver

### Tissue Collection

All animal work was reviewed and approved by the Lawrence Berkeley National Laboratory Animal Welfare and Research Committee or the University of California, Davis Institutional Animal Care and Use Committee.

Mouse fetal tissues were dissected from embryos of different developmental stages from female C57Bl/6N *Mus musculus* animals. Animals, used for obtaining tissue materials from E14.5 and P0 stages, were purchased from both Charles River Laboratories (C57BL/6NCrl strain) and Taconic Biosciences (C57BL/6NTac strain). For tissues of remaining developmental stages, animals (C57BL/6NCrl strain) were purchased from Charles River Laboratories. The number of embryos or P0 pups collected was determined by whether the materials were sufficient for genomic assay, and was not based on statistical considerations. 15-120 embryos/pups were collected for each replicate of each tissue of each stage.

### Tissue Excision And Fixation

See Supplemental File 1-2 for details.

### MethylC-seq Library Construction and Sequencing

MethyC-seq libraries were constructed as previously described^9^ and a detailed protocol is available^64^. An Illumina HiSeq 2500 system was used for all whole genome bisulfite sequencing (WGBS) using either 100 or 130 base single-ended reads.

### Mouse Reference Genome Construction

For all analyses in this study, we used mm10 as the reference genome, which includes 19 autosomes and two sex chromosomes (corresponding to the “mm10-minimal” reference in ENCODE portal, https://www.encodeproject.org/). The fasta files of mm10 were downloaded from UCSC genome browser (Jun 9 2013)^65^.

### WGBS Data Processing

All WGBS data were mapped to the mm10 mouse reference genome as previously described^66^. WGBS processing includes mapping of the bisulfite-treated phage lambda genome spike-in as control to estimate sodium bisulfite non-conversion rate. This pipeline called methylpy is available on github (https://github.com/yupenghe/methylpy). Briefly, cytosines within WGBS reads were first computationally converted to thymines. The converted reads were then aligned by bowtie (1.0.0) onto the forward strand of C-T converted reference genome and the reversed strand of G-A converted reference genome, separately. We filtered out reads that were not uniquely mapped or were mapped to both computationally converted genomes. Next, PCR duplicate reads were removed. Lastly, methylpy counts the methylated basecalls (cytosines) and unmethylated basecalls (thymines) for each cytosine position in the corresponding reference genome sequence (mm10 or lambda).

### Calculation of Methylation Level

Methylation level was computed to measure the intensity and degree of DNA methylation of single cytosines or larger genomic regions. The methylation level is defined as the ratio of the sum of methylated basecall counts over the sum of both methylated and unmethylated basecall counts at one cytosine or across sites in a given region^67^ subtracting the sodium bisulfite non-conversion rate. The sodium bisulfite non-conversion rate is defined as the methylation level of the bisulfite-treated lambda genome.

We calculated this metric for cytosines in both CG context and CH contexts (H=A, C or T). The former is called the CG methylation (mCG) level or mCG level while the latter is called the CH methylation (mCH) level or mCH level.

### ChIP-seq Data Processing

ChIP-seq data were processed using the ENCODE uniform processing pipeline for ChIP-seq. In brief, Illumina reads were first mapped to the mm10 reference using bwa^68^ (version 0.7.10) with parameters “-q 5-l 32-k 2”. Next, the Picard tool (http://broadinstitute.github.io/picard/, version 1.92) was used to remove PCR duplicates using the following parameters: “REMOVE_DUPLICATES=true”.

For each histone modification mark, we represented it as continuous enrichment values of 100bp bins across the genome. The enrichment was defined as the RPKM (Reads Per Kilobase per Million mapped reads) after subtracting ChIP input. The enrichment across the genome was calculated using bamCompare in Deeptools2^69^ using options “–binSize 100–normalizeUsingRPKM–extendReads 300–ratio subtract”. For the ChIP-seq data of EP300, we used MACS^70^ (1.4.2) to call peaks using default parameters.

### RNA-seq Data

Processed RNA-seq data for all fetal tissues, from all stages was downloaded from the ENCODE portal (https://www.encodeproject.org/; Supplemental Table S2).

To further validate our findings regarding transcriptomes generated across laboratories (Wold and Ecker), we generated an additional two replicates of RNA-seq data for fetal forebrain, midbrain, hindbrain and liver tissues. We first extracted total RNA using RNeasy Lipid tissue mini kit from Qiagen (cat no.#74804). Then, we used Truseq Stranded mRNA LT kit (Illumina, RS-122-2101 and RS-122-2102) to constructed stranded RNA-seq libraries on 4ug of the extracted total RNA. An Illumina HiSeq 2500 was used to sequence the libraries and generate 130 bases single-ended reads.

### RNA-seq Data Processing and Gene Expression Quantification

RNA-seq data was processed using the ENCODE RNA-seq uniform processing pipeline. Briefly, RNA-seq reads were mapped to the mm10 mouse reference using STAR^71^ aligner (version 2.4.0k) with Gencode M4 annotation^72^. We quantified gene expression levels using RSEM (version 1.2.23)^73^, expressed as transcripts per million (TPM). For all downstream analyses, we filtered out non-expressed genes and only retained genes that showed non-zero TPM in at least 10% of samples.

### Genomic Features of Mouse Reference Genome

We used GENCODE M4^72^ gene annotation in this study. CG island (CGI) annotation was downloaded from UCSC genome browser (Sep 5, 2016)^65^. CGI shores are defined as the upstream 2kb and downstream 2kb regions along CGIs. Promoters are defined as regions from - 2.5kb to +2.5kb around transcription start sites (TSSs). CGI promoters are defined as those overlapping with CGIs while the remaining promoters are called non-CGI promoters.

We also obtained a list of mappable transposable elements (TEs) using the below procedure. RepeatMasker annotation of the mm10 mouse genome was downloaded from UCSC genome browser (Sep 12, 2016)^65^. The annotation includes 5,138,231 repeats. We acquired the transposon annotation by selecting only the repeats belonging to one of the below repeat classes (repClass): “DNA”, “SINE”, “LTR” or “LINE”. Then, we excluded any repeat elements with a question mark in their name (repName), class (repClass) or family (repFamily). For the remaining 3,643,962 transposons, we further filtered out elements that contained less than 2 CG sites or cases where less than 60% of CG sites within were covered by at least 10 reads across all samples when the data of two replicates were combined. Finally, we utilized the remaining set of 1,688,189 mappable transposable elements for analyses in this study.

### CG Differentially Methylated Region (CG-DMRs)

We identified CG-DMRs using methylpy (https://github.com/yupenghe/methylpy) as previously described^66^. Briefly, we first called CG differentially methylated sites (CG-DMSs) and then merged them into blocks if they both showed similar sample-specific methylation patterns and were within 250bp. Last, we filtered out blocks containing less than three CG-DMSs. In this procedure, we combined the data from the two biological replicates for all tissues, excluding liver samples due to global hypomethylation of the genome.

We overlapped the resulting fetal tissue CG-DMRs with CG-DMRs previously identified in Hon et al^12^ using “intersectBed” from bedtools^74^. The mm9 coordinates of the CG-DMRs from Hon et al. were first, mapped to mm10 using liftOver^65^ with default parameters. Overlap of CG-DMRs is defined as a CG-DMR with at least one base overlap with another CG-DMR when comparing genomic coordinates between lists.

### Identification of Tissue-Specific CG-DMRs

For each fetal tissue type, we defined tissue-specific CG-DMRs as those that showed hypomethylation in a tissue sample from any fetal stage (E10.5 to P0). Hypomethylation is only meaningful with a baseline, thus we used an outlier detection algorithm^75^ to defined the baseline mCG level of each CG-DMR across tissue samples using the mean of the “bulk”, which is defined as the value for the narrowest mCG level interval that includes at least half of all samples. Specifically, 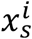 is the mCG level of CG-DMR *i* (*i* = 1, …, *M*) in tissue sample *s* (*s* = 1, …, *N*). Assuming the samples are ordered such that 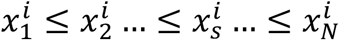, the baseline is defined as the 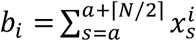 where *a* is the sample index such that 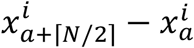 is minimized, i.e. 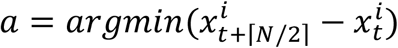 ⌈*N*/2⌉ is defined as the smallest integer that is greater than *N*/2. Lastly, we defined hypomethylated samples as samples in which the mCG level at CG-DMR *i* is at least 0.3 smaller than baseline 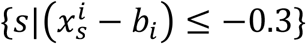. Then, CG-DMR *i* is specific to these tissues. Liver data was not included in this analysis and we excluded CG-DMRs that had zero coverage in any of the non-liver samples. In total, only 402 (~0.02%) CG-DMRs were filtered out.

### Linking CG-DMRs with Genes

We linked CG-DMRs to their putative target genes based on genomic distance. First, we only considered expressed genes, which showed non-zero TPM in at least 10% of all fetal tissue samples. Next, we obtained coordinates for transcriptional start sites (TSSs) of the expressed genes and paired each CG-DMR with the closest TSS using “closestBed” from bedtools^74^. In this way, we inferred a target gene for each CG-DMR; these gene/TSS associations were used in all subsequent analyses in this study.

### Predicting Fetal Enhancer-linked CG-DMRs (feDMRs)

The REPTILE^33^ algorithm was used to identify the CG-DMRs that showed enhancer-like chromatin signatures. We called these fetal enhancer-like CG-DMRs or feDMRs. REPTILE uses a random forest classifier to learn and then distinguish the epigenomic signatures of enhancers and genomic background. One unique feature of REPTILE is that by incorporating the data of additional samples (as outgroup/reference), it is able to employ epigenomic variation information to improve enhancer prediction. In this study, REPTILE was run using input data from CG methylation (mCG) and six histone marks (H3K4me1, H3K4me2, H3K4me3, H3K27ac, H3K27me3 and H3K9ac).

A REPTILE enhancer model was trained as previously described^33^. Briefly, CG-DMRs were called across the methylomes of mouse embryonic stem cells (mESCs) and all eight E11.5 mouse tissues. CG-DMRs were required to contain at least 2 CG-DMSs and they were extended 150bp in each direction (5’ and 3’). The REPTILE model was trained on the mESC data using E11.5 mouse tissues as outgroup. Data from mCG and six histone modifications are available for these samples. The training dataset consists of 5000 positive instances (putative known enhancers) and 35,000 negative instances. Positives were 2kb regions centered at the summits of top 5,000 EP300 peaks in mESCs. Negatives include randomly chosen 5,000 promoters and 30,000 2kb genomic bins. The bins have no overlap with any positives or promoters. REPTILE learned the chromatin signatures that distinguish positive instances from negative instances.

Next, using this enhancer model, we applied REPTILE to delineate feDMRs from the 1,808,810 CG-DMRs identified across all non-liver tissues. feDMRs were predicted for each sample based on data from mCG and six core histone marks, while the remaining non-liver samples were used as an outgroup. In REPTILE, the random forest classifier for CG-DMR assigns a confidence score ranging from 0.0 to 1.0 to each CG-DMR in each sample. This score corresponds to the fraction of decision trees in the random forest model that vote in favor of the CG-DMR to be an enhancer. Previous benchmarks showed that the higher the score, the more likely that a CG-DMR shows enhancer activity^33^. We named this confidence score as the enhancer score. For each tissue sample, feDMRs are defined as CG-DMRs with enhancer score greater than 0.3. feDMRs were also defined for each tissue type as the CG-DMRs that were identified as an feDMR in at least one tissue sample of that tissue type. For example, if a CG-DMR was predicted as an feDMR only in E14.5 forebrain, it was classified as a forebrain-specific feDMR.

We overlapped the feDMRs with putative enhancers from Yue et al^37^. We utilized a set of coordinates identifying the center base position of putative enhancers for each of the tissues and cell types from http://yuelab.org/mouseENCODE/predicted_enhancer_mouse.tar.gz. Next, we defined putative enhancers as +/-1kb regions around the centers. Putative enhancers from different tissues and cells types were combined and merged if they overlapped. The merged putative enhancers (mm9) were then mapped to the mm10 reference using liftOver^65^. Finally, “intersectBed” from bedtools^74^ were used to overlap feDMRs with these putative enhancers.

### Enhancer Score and *in vivo* Enhancer Activity

To estimate the likelihood that an feDMR with a certain enhancer score actually displays enhancer activity *in vivo*, we utilized enhancer data set from the VISTA enhancer browser^38^. Using the DNA elements in VISTA enhancer browser (VISTA elements), we validated feDMRs identified in six E11.5 tissues (forebrain, midbrain, hindbrain, heart, limb and neural tube), where at least 30 validated VISTA elements (enhancers) are available. Specifically, for each E11.5 tissue, we first overlapped feDMRs predicted in that tissue with VISTA elements and identified VISTA elements that fully contained at least one feDMR. Then, we calculated the fraction of feDMR overlapping VISTA elements that displayed enhancer activity in the predicted tissue, any tissue(s) as well as those that did not display enhancer activity in any tissue.

### Enriched Transcription Factor (TF) Binding Motifs in Tissue-Specific feDMRs

To identify TF motifs enriched in feDMRs, we scanned the genome to delineate TF motif occurrences as previously described^46^. Briefly, we utilized TF binding position weight matrices (PWMs) from the MEME motif database (v11, 2014 Jan 23. motif sets chen2008, hallikas2006, homeodomain, JASPAR_CORE_2014_vertebrates, jolma2010, jolma2013, macisaac_theme.v1, uniprobe_mouse, wei2010_mouse_mw, wei2010_mouse_pbm, zhao2011). Then, FIMO^76^ was used to scanned the genome to identify TF motif occurrences using options “–output-pthresh 1E-5–max-stored-scores 500000”.

Next, we performed a hypergeometric test to identify significant motif enrichments. For each tissue type, we calculated the motif enrichment for feDMRs in that tissue (foreground) against a list of feDMRs identified for other tissues not overlapping with the foreground tissue list. For this analysis, we extended the average size of both foreground and background feDMRs to 400bp to avoid bias due to size differences. For a given tissue *t*, the total number of foreground and background feDMRs is *N*_*f,t*_ and *N*_*b,t*_, respectively, and *N*_*t*_ = *N*_*f,t*_ + *N*_*b,t*_ is the total number of feDMRs. For a given TF binding motif *m*, TF motif occurrences are overlapped with *n*_*f,t,m*_ foreground and *n*_*b,t,m*_ background feDMRs, while *n*_*t,m*_ = *n*_*f,t,m*_ + *n*_*b,t,m*_ is the total number of overlapping feDMRs. The probability of observing *n*_*f,t,m*_ or more overlapping foreground feDMRs (p-value) is defined as:

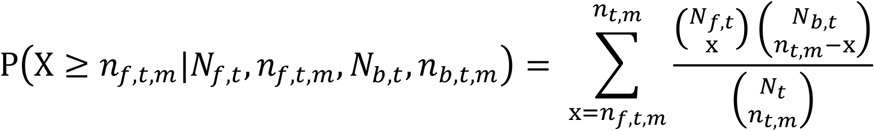

For each tissue type, we performed this test for all motifs (n=532). Then, the p-values of each tissue were adjusted using Benjamini-Hochberg method and the motifs were called as significant if they passed 1% FDR cutoff. Lastly, we excluded any TF-binding motifs whose TF expression level was less than 10 TPM.

### Enrichment of GWAS SNPs in feDMRs

A set of genome-wide association study (GWAS) SNPs were utilized from the GWAS Catalog^77^ (gwas_catalog_v1.0-associations_e86_r2016-11-28.tsv). We filtered out SNPs with missing coordinates or missing p-value information as well as those with p-values greater than 5x10^-8^, resulting in a final set of 13,470 GWAS SNPs. Next, we used liftOver from the UCSC genome browser^65^ to convert the SNP coordinates from hg38 human reference genome to hg19, resulting in the exclusion of only 4 SNPs. We further selected SNPs that could be lifted over to mm10 mouse reference genome, resulting in a final set of 7,052 GWAS SNPs that are conserved between human and mouse; these were included in the following analysis.

Next, to obtain orthologous regions from the human genome of the mouse CG-DMRs, we used liftOver to map mouse CG-DMRs (mm10) to hg19, requiring that at least 50% of the bases in CG-DMR could be assigned to hg19 (using option - minMatch=0.5). In total, 1,034,801 out of 1880810 (55%) of mouse DMR regions could be aligned to the human genome.

Finally, we overlapped the human orthologous regions of the mouse feDMRs for each tissue/organ with the GWAS SNPs and tested for enrichment using a one-tailed hypergeometric test. Specifically, for each tissue/organ, we separately overlapped GWAS SNPs with the human orthologous regions for distal feDMRs in that tissue/organ (foreground) and the human orthologous regions for the remaining CG-DMRs (background), Using the statistical approach (described below) for SNPs associated with each trait, we tested for their enrichment in the human orthologous regions for distal feDMRs compared to background. We then used Benjamini-Hochberg approach to adjust the p-values for multiple testing. P-value cutoff given 5% false discovery rate (FDR) was used to call significant enrichment. This procedure was conducted separately for the distal feDMRs of each tissue/organ.

In brief, we used a one-tailed hypergeometric test to test for enrichment. We defined q_*c*_ as the number of GWAS SNPs that are associated with trait c and are overlapped with distal feDMRs. Therefore, the total number of SNPs overlapped with foreground is Q =∑_c_ q_c_. Similarly, we defined B =∑_c_ b_c_ as the number of SNPs that are overlapped with background where b_c_ SNPs were associated with trait *c*. For each trait c, the null hypothesis was that q_c_ follows a hypergeometric distribution, with population size as *N* = *Q* + *B*, number of successes (here the number of SNPs related to *c* that are overlapped with either foreground or background) as n_c_ = *q*_c_ + *b*_c_, and sampling number as *Q*. Let X be a random variable representing the observed number of SNPs that are related to trait *c* and overlapped with foreground. Thus, the probability of observing X is equal to or greater than *q*_c_ (i.e. p-value) is calculated as:

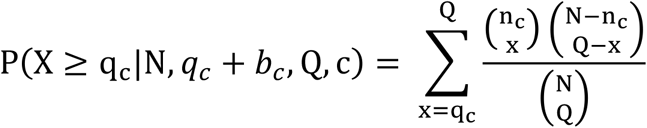

### Categorizing CG-DMRs

To better understand the potential functions of CG-DMRs, we grouped them into various categories based on their genomic location and chromatin signatures. First, we overlapped CG-DMRs with promoters, CGIs and CGI shores and defined the CG-DMRs overlapping these locations as proximal CG-DMRs. Out of the 153,019 proximal CG-DMRs, 46,692, 90,831, 1,710 and 13,786 overlapped with CGI promoters, non-CGI promoters, CGIs and CGI shores, respectively. We avoided assigning proximal CG-DMRs into multiple categories by prioritizing the four genomic features as CGI promoter, non-CGI promoter, CGI and CGI shores (ordered in decreasing priority). Each CG-DMRs was assigned to the category with the highest priority.

We further classified the remaining 1,655,791 distal CG-DMRs: 1) 415,227 of them were predicted as distal feDMRs (CG-DMRs that show enhancer-like chromatin signatures^34,35^) as described in the previous method section. (2) Next, we defined flanking distal feDMRs or fd-feDMRs as the CG-that were within 1kb to distal feDMRs but were not predicted as enhancers (feDMRs). In total we found 221,960 such CG-DMRs.

(3) Then, among the remaining, unclassified CG-DMRs, 194,610 CG-DMRs were identified tissue-specific CG-DMRs in at least one of the tissues because they displayed strong tissue-specific hypomethylation patterns (mCG difference ≥ 0.3). By checking the enrichment of histone marks in their hypomethylated tissues, we found they were enriched for H3K4me1 but not other histone marks, and such chromatin signatures resembled that of primed enhancers^42^. Therefore, we defined these CG-DMRs as primed distal feDMRs or pd-feDMRs.

(4) Lastly, we defined the remaining CG-DMRs as unexplained CG-DMRs (unxDMRs) because their functional roles could not be assigned yet. We found unxDMRs have strong overlap with transposable elements and we further divided them into two classes: te-unxDMRs and nte-unxDMRs. te-unxDMRs are unxDMRs that are overlapped with transposable elements, while the remaining were nte-unxDMRs.

### Evolutionary Conservation of CG-DMRs

The evolutionary conservation of CG-DMRs were measured using phyloP score^78^ from the UCSC genome browser^65^ (http://hgdownload.cse.ucsc.edu/goldenpath/mm10/phyloP60way/mm10.60way.phyloP60way.bw). Next, Deeptools2^69^ was used to generate the profile of evolutionary conservation of the CG-DMR centers and +/-5kb flanking regions using options “reference-point–referencePoint=center -a 5000 -b 5000”.

### Finding TF-binding Motifs Enriched in Flanking Distal feDMRs

To identify the TF-binding motifs enriched in fd-feDMRs relative to feDMRs, we performed motif analysis using the former as foreground and the latter as background. Specifically, for each tissue, the tissue-specific feDMRs were used as background, while fd-feDMRs that were within 1kb of these tissue-specific feDMRs were used as foreground. To avoid potential bias residing in different size distribution, both foreground and background regions were extended from both sides (5’ and 3’) such that both had mean size of 400bp. Next a hypergeometric test was performed to find TF-binding motifs that were significantly enriched in foreground. This test was the same as that used for the identification of TF-binding motifs in feDMRs.

### TF-binding Motif Enrichment Analysis for Primed Distal feDMRs

We also performed motif analysis to identify TF-binding motifs enriched in pd-feDMRs. The procedure was similar to the motif enrichment analysis on feDMRs. For each tissue, the pd-feDMRs hypomethylated in that tissue were considered as foreground while the remaining pd-feDMRs were considered as background. Then, a hypergeometric test was performed to identify significant motif enrichment.

Next, for each tissue type, we compared the TF-binding motifs enriched in pd-feDMRs and the tissue-specific feDMRs. The hypergeometric test was used to test the significance of overlap – the chance of obtaining the observed overlap if the two lists were based on random sampling (without replacement) from the TF-binding motifs with TF expression level greater than 10 TPM.

### Permutation Test of the Overlap between unxDMR and TEs

To estimate the significance of overlap between unxDMRs and TEs, we shuffled the location of unxDMRs using “shuffleBed” tool from bedtools^74^ with default setting and recalculated the overlaps. After repeating this step for 1,000 times, we obtained an empirical estimate of the overlap if unxDMRs were randomly distributed in the genome. Let the observed number of TE overlapping unxDMRs be *x*^*obs*^ and the number of TE overlapping shuffled unxDMRs in permutation *i* be 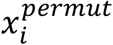. We then calculated p-values as

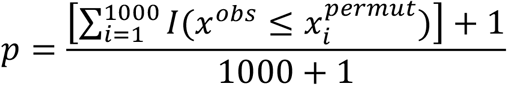

where 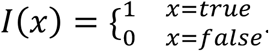.

### Quantification of the mCG Dynamics in Tissue-Specific CG-DMRs

To quantify mCG dynamics, we defined and counted loss-of-mCG and gain-of-mCG events. A loss-of-mCG (Gain-of-mCG) event is a decrease (increase) of mCG level by at least 0.1 in one CG-DMR in one stage interval. For example, if the mCG level of one CG-DMR at E11.5 and E12.5 is 0.8 and 0.7 in heart respectively, it is considered a loss-of-mCG event occurred at the stage interval E11.5-E12.5. Stage interval is defined as the transition between two sampled adjacent stages(e.g. E15.5 and E16.5).

### Clustering Forebrain-specific CG-DMRs based on mCG and H3K27ac Dynamics

We used k-means clustering to identify subgroups of forebrain-specific CG-DMR based on mCG and H3K27ac dynamics. First, for each forebrain-specific CG-DMR, we calculated the mCG level and H3K27ac enrichment in forebrain samples from E10.5 to adult stages. Here, we used methylome data for postnatal 1, 2 and 6 week frontal cortex from Lister et al^10^ to approximate the DNA methylation landscape of adult forebrain. We also incorporated H3K27ac data for postnatal 1, 3 and 7 week forebrain samples. Next, to make the range H3K27ac enrichment values comparable to that of mCH levels, for each forebrain-specific CG-DMR, the negative H3K27ac enrichment values were thresholded as zero and then each value was divided by the maximum. If the maximum was zero for some forebrain-specific CG-DMRs, we set all values to be zero. k-means clustering was of subgroups was carried out but no new patterns were observed. Lastly, we used GREAT^79^ employing the “Single nearest gene” association strategy to identify the enriched gene ontology terms of genes near CG-DMRs for each subgroup.

### Association Between mCG level and H3K27ac Enrichment

To investigate the association between mCG and H3K27ac, for each tissue and each developmental stage, we first divided the tissue-specific CG-DMRs into three categories based on mCG methylation levels: H (highly CG methylated; mCG level > 0.6), M (moderately CG methylated; 0.2 < mCG level ≤ 0.6) and L (lowly CG methylated; mCG level ≤ 0.2). Then, we examined the distribution of H3K27ac enrichment in different groups of CG-DMRs by counting the number of CG-DMRs for each of four levels of H3K27ac: [0,2], (2, 4], (4, 6] and (6, ∞).

### Weighted Correlation Network Analysis (WGCNA)

We used weighted correlation network analysis (WGCNA)^80^, an unsupervised method, to detect sets of genes with similar expression profiles across samples (R package, “WGCNA” version 1.51). Briefly, TPM values were First log2 transformed (with pseudo count 1e-5). Then, the TPM value of every gene across all samples was compared against the expression profile of all other genes and a correlation matrix was obtained. To obtain connection strengths between any two genes, we transformed this matrix to an adjacency matrix using a power adjacency function. To choose the parameter (soft threshold) of the power adjacency function, we used the scale-free topology (SFT) criterion, where the constructed network is required to at least approximate scale-free topology. The SFT criterion recommends use of the first threshold parameter value where model-fit saturation is reached as long as it is above 0.8. In this study, the threshold was reached for a power of 5.

Next, the adjacency matrix is further transformed to a topological overlap matrix (TOM) that finds “neighborhoods” of every gene iteratively, based on the connection strengths. The TOM was calculated based on the adjacency matrix derived using the signed hybrid network type, biweight mid correlation and signed TOMtype parameters of the TOMsimilarityFromExpr module in WGCNA. Hierarchical clustering of the TOM was done using the flashClust module using the average method. Next, we used the cutreeDynamic module with the hybrid method, deepSplit = 3 and minClusterSize = 30 parameters to identify modules that have at least 30 genes. A summarized module-specific expression profile was created using the expression of genes within the given module, represented by the eigengene. The eigengene is defined as the first principal component of the log2 transformed TPM values of all genes in a module. In other words, this is a virtual gene that represents the expression profile of all genes in a given module. Next, very similar modules were merged after a hierarchical clustering of the eigengenes of all modules applying a distance threshold of 0.15. Finally, the eigengenes were recalculated for all modules after merging.

### Gene Ontology Analysis of Genes in Co-expression Modules (CEM)

To better understand the biological processes of genes in each CEM, we used Enrichr^81,82^ (http://amp.pharm.mssm.edu/Enrichr/) to identify the enriched gene ontology terms in the “GO_Biological_Process_2015” category.

### Correlating Eigengene Expression with mCG and Enhancer Scores of feDMRs

We investigated the association between gene expression and epigenomic signatures of regulatory elements in CEMs. First, for each CEM, we used the eigengene expression to summarize the transcription patterns of all genes in the module. Then, we calculated the normalized average enhancer score and normalized average mCG level of all feDMRs that were linked to the genes in the CEM. Specifically, to reduce the potential batch effect, for each tissue and each stage, we normalized the enhancer score of each feDMR by the mean enhancer score of all feDMRs. mCG levels of feDMRs were normalized in similar way except that the data of all DMRs was used to calculate the mean mCG level for each tissue and each stage. Next, for each CEM, the TPM of its eigengene, the normalized average enhancer score and mCG level of linked feDMRs were converted to z-scores across all fetal stages for each tissue type (for analysis for tissue-specific expression) or across tissue types for each development stage (for analysis for temporal expression). Lastly, for each CEM, we calculated the Pearson correlation coefficient (R 3.3.1) between the z-score of eigengene expression and the z-score of normalized enhancer score (or mCG level) for each module. The correlation coefficients were calculated for two different settings: 1) for each tissue type, the correlation was computed using z-score of normalized eigengene expression values and enhancer scores (or mCG levels) across different development stages or 2) for each developmental stage, the correlation was computed across different tissue types. The coefficients from the former analysis indicate how well temporal gene expression is correlated with enhancer score or mCG level of regulatory elements, while the latter measures the association with tissue-specific gene expression.

We then test whether the correlation that we observed was significant by comparing it with the correlation based on shuffle data. In the analysis for tissue-specific expression in a given tissue type, we mapped the eigengene expression of one CEM to the enhancer score (or mCG level) of feDMRs linked to genes in a randomly chosen CEM. For example, in the shuffle setting, when given tissue type was heart, we calculated the correlation between the eigengene expression of CEM14 and the enhancer score of the feDMRs linked to genes in CEM6. In the analysis for temporal expression, given a specific developmental stage, we performed similar permutation. Next, we calculated the Pearson correlation coefficients for this permutation setting. Lastly, using a two-tailed Mann-Whitney test, we compared the median of observed correlation coefficients and the median of those based on shuffled data.

### Identification of Large Hypo CG-DMRs

Large hypo CG-DMRs were called using the same procedure as previously described^46^. For each tissue type, tissue-specific CG-DMRs were merged if they are within 1kb of each other. Then, we filtered out merged CG-DMRs less than 2kb in length.

We overlapped genes with large hypo CG-DMRs and then filtered out any genes with names starting with “Rik” or “Gm[0-9]”, where [0-9] represents a single digit, because the ontology of these genes were ill-defined.

### Super-enhancer Calling

Super-enhancers were identified using ROSE^83,84^ pipeline. First, H3K27ac peaks were called using macs2^70^ callpeak module with options “–extsize 300 -q 0.05–nomodel-g mm”. Control data was used in the peak-calling step. Next, ROSE was run with options “-s 12500 -t 2500”, and H3K27ac peaks, mapped H3K27ac ChIP-seq reads and mapped control reads as input. The super-enhancers calls were generated for each tissue sample. Then, we obtained the super-enhancers of one tissue type by merging the super-enhancers called at each stage of fetal development (E10.5 to P0). Last, we generated a list of merged super-enhancers by merging super-enhancer calls of all tissue types except liver.

### DNA Methylation Valley (DMV) Identification

We identified DMVs as previously described^55^. First, genome was dividing into 1kb non-overlapping bins. Then, for each tissue sample (replicate), consecutive bins with mCG level less than 0.15 were merged into blocks; bins with no data (no CG sites or no reads) were skipped. Next, any blocks merged from no less than 5 with-data bins were called as DMVs. For each tissue sample, we filtered for DMVs reproducible in two replicates by first selecting the DMVs identified in one replicate that are overlapped any DMVs called in the other replicate, and then merging overlapping DMVs. Using this strategy, we obtained DMV calls for each tissue from each developmental stage. Lastly, we got a list of merged DMVs for all tissue samples by merging all DMVs identified in any tissues from any developmental stages.

We overlapped genes with DMVs and then filtered out any genes with names starting with “Rik” or “Gm[0-9]”, where [0-9] represents a single digit, because the ontology of these genes were ill-defined.

### Partially Methylated Domain (PMD) Identification

PMDs were identified as previously described^9^ using random forest classifier. To train the classifier, we first visually selected regions on chromosome 19 as strong candidates for PMDs or non-PMDs in E14.5 liver sample. Specifically, we manually annotated 5 PMDs with which showed obvious lower mCG level compared to adjacent genomic regions (chr19:46110000-46240000, chr19:45820000-45960000, chr19:47140000-47340000 and chr19:48060000-52910000) and 7 Non-PMD regions (chr19:4713800-4928700, chr19:7420700-7541100, chr19:8738100-8967000, chr19:18633300-18713800, chr19:53315500-53390000, chr19:55256600-55633900 and chr19:59281600-59329200).

Next, these regions were divided into 10kb non-overlapping bins and we calculated the percentiles of the methylation levels at the CG sites within each bin. CG sites that were within CGIs, DMVs^55^ or any of four Hox loci (see below) were excluded as these regions are typically hypomethylated which may result in incorrect PMD calling. Additionally, sites with less than 5 reads covered were also excluded. We trained the random forest classifier using data from E14.5 liver (combining the two replicates) and we then predicted whether a 10kb bin was a PMD or non-PMD in all liver samples (considering replicates separately). We chose a large bin size (10 kb) to reduce the effect of smaller scale methylation variation (such as DMRs) as PMDs were first discovered as large (mean length = 153kb, PMID: 19829295) regions with intermediate methylation level (< 70%, PMID: 19829295). Furthermore, the features (the distribution of methylation level of CG sites, which measured the fraction of CG sites that showed methylation level at various methylation level ranges) used in the classifier required enough CG sites within each bin to robustly estimate the distribution, which necessitated a relatively large bin. Also, we excluded any 10kb bins containing less than 10 CG sites for the same reason. These percentiles were used as features for the random forest. The random forest implement was from scikit-learn (version 0.17.1)^85^ python module and the following arguments were supplied to the Python function RandomForestClassifier from scikit-learn: n_estimators = 10000, max_features=None, oob_score=True, compute_importances=True.

Lastly, we merged consecutive 10kb bins that were predicted as PMD into blocks and filtered out blocks smaller than 100kb. We further excluded blocks that overlapped with gaps in mm10 genome (downloaded from UCSC genome browser, Sep 21, 2013). To obtain a set of PMDs that were reproducible in both replicates, we only considered genomic regions that were larger than 100kb and were covered by PMD calls in both replicates. These regions were the final set of PMDs used for later analyses. Because there was only one replicated for adult liver, we called the PMDs at this stage using the single replicate.

PMDs were originally called using the above procedure without excluding CG sites in Hox gene clusters. However, because these Hox loci are more likely to be considered as large DMVs^55^, we removed any PMD that overlapped with the four Hox clusters (chr11:96257739-96358516, chr15:102896908-103038064, chr2:74648392-74748841 and chr6:52146273-52277140).

### Overlap Between PMDs and LADs

To examine the relationship between PMDs and lamina associated domains (LADs) in normal mouse liver cells (AML12 hepatocyte) we utilized LAD data in the Table S2 from Fu et al^86^. The mm9 coordinates of LADs were converted to mm10 using liftOver with default settings. We then used permutation testing to examine the significance of the overlap between PMDs and LADs. Similar to the procedure for checking the overlap between TEs and unxDMRs, we permutated (1000 times) the genomic locations of PMDs and recorded the number of overlapping bases (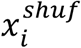 for permutation *i*) between shuffled PMDs and LADs. Then, we compared 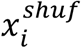 with the observed numbers of overlapping bases (*x*^*obs*^) between PMDs and LADs and computed p-values as:

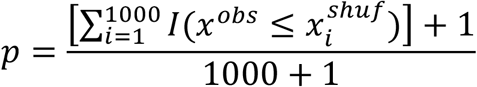

where 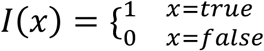.

### Replication Timing Data

Replication timing data (build mm10) of three mouse cell types was utilized from ReplicationDomain^87^. The cell types used for these analyses were mESC (id: 1967902&4177902_TT2ESMockCGHRT), neural progenitor cells (id: 4180202&4181802_TT2NSMockCGHRT) and mouse embryonic fibroblasts (id: 304067-1 Tc1A).

### Gene Expression in PMDs

We obtained PMD overlapping protein-coding genes information using “intersectBed”. A similar approach was used to obtain the protein-coding genes overlapped with PMD flanking regions (upstream 100kb and downstream 100kb of PMDs); genes overlapping with PMDs were removed from this list. Lastly, we compared the expression of PMD-overlapping genes (n=5,748) and the genes (n=2,555) overlapping with flanking regions.

### Sequence Context Preference of mCH

To interrogate the sequence preference of mCH, as previous described^9^, we first identified CH sites that showed a significantly higher methylation level than the low level noise (which was around 0.005 in term of methylation level) caused by incomplete sodium bisulfite non-conversion. For each CH site, we counted the number of reads that supported methylation and the number of reads that did not. Next, we performed a binomial test with the success probability equal to the sodium bisulfite non-conversion rate. FDR (1%) was controlled using the Benjamini-Hochberg approach^88^. This analysis was independently performed for each three-nucleotide context (e.g., a pvalue cutoff was calculated for CAG cytosines). Lastly, we counted sequence motif occurrence of +/-5bp around the tri-nucleotide context of methylated mCH sites and visualized the sequence preferences using seqLogo^89^

### Calling mCH Domains

We used an iterative process to call mCH domains, which are genomic regions that are enriched for mCH compared to flanking regions. First, we selected a set of samples that showed no evidence of mCH. Data from these samples were used in the following steps to filter out genomic regions that are prone to misalignment and showed suspicious mCH abundance. Analysis of the global mCH level and mCH motifs revealed that E10.5 and E11.5 tissues (excluding heart samples) have extremely low mCH and the significantly methylated non-CG sites showed little CA preference. Therefore, we assumed these sites contain no mCH domain and any mCH domain called in control samples by the algorithm were likely artifacts. By filtering out the domains called in the control samples, we were able to exclude the genomic regions that were prone to mapping error or avoid other potential drawbacks in the processing pipeline.

In order to identify genomic regions where sharp changes in mCH levels occurred, we applied a change point detection algorithm with the mCH levels of all 5kb non-overlapping bins across the genome as input. We only included bins that contained at a minimum 500 CH sites and at least 50% of CH sites were covered by 10 or more reads. The identified regions defined the boundaries that separate mCH domains from genomic regions showing background level mCH. We implemented this step using the function *cpt.mean* in R package “changepoint”, with options “method="PELT", pen.value=0.05, penalty="Asymptotic" and minseglen=2”. To match the range of chosen penalty, we scaled up mCH levels by a factor of 1,000.

The iterative procedure was carried out as follows: 1) An empty list of excluded regions was created. 2) For each control sample, the change point detection algorithm was applied to the scaled mCH levels of 5kb non-overlapping bins. Bins overlapping with excluded regions were ignored. 3) The genome was segmented into chunks based on identified change points. 4) The mCH level of each chunk was calculated as the mean mCH level of the overlapping 5kb bins that were not overlapped with excluded regions. 5) mCH domains were identified as chunks whose mCH level was at least 50% greater than the mCH level of both upstream and downstream chunks. The pseudo mCH level of 0.001 was used to avoid dividing zero. 6) mCH domains were added to the list of excluded regions. 7) Step 2 to 6 were repeated until the list of excluded regions stop expanding. 8) Steps 2-5 were then applied to all samples. 9) For each tissue/organ, only regions were retained that were identified as (part of) an mCH domain in both replicates and regions less than 15kb in length were filtered out; mCH domains must span at least three bins. The above criterion were used to define mCH domains for each tissue/organ. 10) Individual mCH domains from each tissue and organ were merged to obtain a single combined list of 384 mCH domains.

### Clustering of mCH Domains

We applied k-means clustering to group the 384 identified mCH domains into 5 clusters based on the normalized mCH accumulation profile of each mCH domain and corresponding flanking regions (100kb upstream and 100kb downstream). Specifically, 1) in each tissue sample, the mCH accumulation profile of one mCH domain was represented as a vector of length 50: the mCH level of 20 5kb bins upstream mCH domain, 10 bins that equally divided the mCH domain and 20 5kb bins downstream. 2) Then, we normalized all values by the average mCH level of bins of flanking regions (the 20 5kb bins upstream and 20 5kb bins downstream of mCH domain). 3) We next computed the profile in samples of 6 tissue types (midbrain, hindbrain, heart, intestine, stomach and kidney) that showed the most evident mCH accumulation in fetal development. 4) Using the profile of these tissue samples, k-means (R v3.3.1) was used to clustered mCH domains with k = 5. We also tried higher cluster numbers (e.g. 6) but did not identify any new patterns. Even using the current k setting (k=5), the mCH domains in clusters 1 (C1) and 3 (C3) shared similar mCH accumulation pattern.

### Genes in mCH Domains

We obtained the overlapping gene information for each of the mCH domains by overlapping gene bodies with mCH domains using “intersectBed” in bedtools^74^. Only protein coding genes were considered. We further filtered out any genes with names starting with “Rik” or “Gm[0-9]”, where [0-9] represents a single digit, because the ontology of these genes were ill-defined. For the overlapping genes of each mCH domain cluster, we used EnrichR^81,82^ to find the enriched gene ontology terms (“GO_Biological_Process_2015”).

Next we asked whether the identified overlapping genes were enriched for TF encoding genes. For this purpose, a list of mouse TFs from AnimalTFDB^90^ (Feb 27, 2017) was utilized. We then performed a permutation test to estimate the significance of the findings. Specifically, *x*^*obs*^ is the number of TF encoding genes in all overlapping genes. We randomly selected (1000 times) the same number of genes and in the *i*th time, 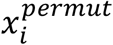 of the randomly selected genes encoding TFs. Lastly, the p-values was calculated as

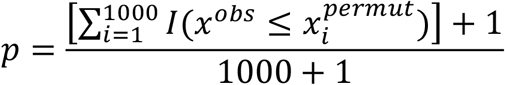

where 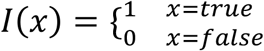.

### mCH Accumulation Indicates Gene Repression

To evaluate the association between mCH abundance and gene expression, we traced the expression dynamics of genes inside mCH domains. For mCH domains in each cluster, we first calculated the TPM z-score for each of the overlapping genes. Specifically, for each tissue type and each overlapping gene, we normalized TPM values in the samples of that tissue type to z-scores. The z-scores showed the trajectory of dynamic expression, in which the aptitude information of expression was removed. If the gene was not expressed, we did not perform the normalization. Next, we calculated the z-scores for all genes that had no overlapped with any mCH domain. Lastly, we subtracted the z-scores of overlapping genes by the z-scores of all genes outside mCH domains. The resulting values indicated the level of expression of genes in mCH domains relative to genes not in mCH domains.

### feDMRs in mCH Domains

To evaluate whether feDMRs were enriched in mCH domains, we calculated the percentage of bases within mCH domains that were also within tissue-specific feDMRs. Specifically, we first divided the genome into non-overlapping 100bp bins and then, for each tissue, we calculated the percentage of bases in each bin that overlapped with tissue-specific feDMRs. Next, we plotted such percentages across mCH domains (each was equally divided into 10 non-overlapping bins) and flanking regions (10kb upstream and 10kb downstream; each contained 10 1kb bins).

For the feDMRs that overlapped with mCH domains, we then traced their mCG levels as they changed over developmental time. For each tissue type and each developmental stage, we calculated the percentage of tissue-specific feDMRs whose mCG level was in each range: [0-0.2), [0.2, 0.4), [0.4, 0.6), [0.6, 0.8) and [0.8, 1]. For each tissue and each developmental stage, these percentages were calculated for tissue-specific feDMRs in mCH domains and also for all tissue-specific feDMRs. Lastly, the ratio of the former to the later was defined as the enrichment of feDMRs (with certain mCG level) in mCH domains, which was then transformed to log_2_ ratio.

